# The penta-EF-hand protein Pef1 of *Candida albicans* functions at sites of membrane perturbation to support polarized growth and membrane integrity

**DOI:** 10.1101/2024.09.06.611525

**Authors:** Martin Weichert, Marcel René Schumann, Ulrike Brandt, Alexandra C. Brand, André Fleißner

**Affiliations:** Institut für Genetik, Technische Universität Braunschweig, Braunschweig, Germany; Medical Research Council Centre for Medical Mycology (MRC CMM) at the University of Exeter, Exeter, United Kingdom

**Keywords:** <u>Keywords</u>: *Candida albicans*, Cell membrane integrity, membrane stress, membrane repair, polyene, saponin, cell polarity, polarized growth, PEF-hand protein, Pef1, calcium, calcineurin, pathogenicity

## Abstract

The fungal plasma membrane is the target of fungicidal compounds, such as polyenes and saponins, that directly interact with fungus-specific ergosterol to cause deleterious membrane disruption. To counter membrane attack, diverse eukaryotic cells employ Ca^2+^-binding penta-EF (PEF)-hand proteins, including the human ortholog, ALG-2, to maintain membrane integrity. *Candida albicans* is a major fungal pathogen in humans, where increasing resistance to current antifungal drugs that target the plasma membrane is a serious cause of concern. Combinatorial treatments that additionally compromise the plasma membrane offer a way forward, but our mechanistic understanding of how fungi respond to direct membrane disruption remains limited. Here, we characterized the PEF-hand ortholog, Pef1, in this polymorphic species. GFP-tagged Pef1 localized at sites of polarized growth in yeast and hyphal cells of *C. albicans*. On treatment of cells with the polyene drug, amphotericin B, or the saponin, tomatine, GFP-Pef1 appeared as punctate spots at the membrane. In a similar manner, loss of calcineurin, but not of its transcription factor, Crz1, caused a punctate localization pattern of GFP-Pef1, which correlated with the serum sensitivity of the *cna1*Δ mutant. While deletion of *PEF1* impaired yeast cell separation, filamentation was not affected. Strikingly, *pef1*Δ hyphae could not maintain plasma membrane integrity in serum. Consistent with this, the mutant exhibited attenuated virulence in an insect larvae infection model. Taken together, these observations suggest that Pef1 localizes to sites of membrane perturbation in order to maintain cell integrity, including sites of dynamic polarized growth and fungicide-induced membrane disruption.

## Introduction

Maintaining and promoting the integrity and function of cell membranes is vital for all eukaryotic organisms, including fungi. Although fungal cells are surrounded by dynamic cell walls that provide major protection from mechanical, chemical or osmotic insults (1), the underlying plasma membrane can still be exposed to various forms of stress that pose a major threat to cellular integrity and survival. Fungal lipid bilayers contain ergosterol, a sterol that is fungus-specific and therefore an attractive target in the search for compounds that directly interact with it (2, 3). Ergosterol is crucial for maintaining both the barrier function of the plasma membrane and the functionality of membrane-associated and transmembrane proteins that support cell wall structure, nutrition, cell polarity and stress signaling pathways of fungal cells (4, 5). Ergosterol-binding antifungal metabolites from the polyene and saponin classes, which are unrelated fungicides naturally produced by bacteria and plants, quickly disrupt fungal cell membrane integrity via ergosterol extraction or deleterious membrane-pore formation (6, 7). Because of their fungicidal activity, nystatin and amphotericin B are used as polyene drugs in the treatment of infectious disease caused by human-pathogenic fungi, including *Aspergillus fumigatus* and *Candida albicans* (8, 9). Infections of plants caused by phytopathogenic fungi such as *Botrytis cinerea* are often followed by the release of saponins, for example α-tomatine from tomato plants, sparking interest in understanding how saponins could be exploited to combat fungal infections and to boost plant defenses (10, 11). To combat pathogenic fungi in clinical and agricultural settings, the azole class of antifungals is widely employed to block ergosterol biosynthesis, but their indirect and relatively slow effect on fungal membranes is mostly fungistatic and can select for antifungal resistance, threatening the applicability of azoles (2). Although fungicides, as well as membrane-disrupting antifungal immune responses in humans, animals and plants, are highly relevant and efficient in directly targeting fungal membranes (12, 13), our current knowledge about the molecular basis of the membrane stress response in fungi during attack and defense remains limited.

The general membrane stress response in fungi involves the activation of important pathways, including the well-studied Ca^2+^/calcineurin-dependent transcriptional response (14, 15). However, this signaling pathway does not explain how fungi immediately respond to direct and acute forms of membrane disruption. In light of this, our previous study in the model fungus, *Neurospora crassa*, identified a role for the penta-EF (PEF)-hand protein, PEF1, in mediating a quick response to deleterious membrane disruption that promotes cellular survival (16). Pef1 belongs to a class of cytosolic eukaryotic proteins that feature five highly-conserved Ca^2+^-binding helix-loop-helix motifs (17). In *N. crassa*, the PEF1 protein facilitates membrane integrity during plasma membrane fusion of interacting vegetative cells. Strikingly, PEF1 is recruited to the ergosterol-rich polarized cell tips of filaments during the exposure to nystatin or tomatine, and this protein directly contributes to the tolerance of the saponin (16). This protective role against tomatine was also demonstrated in *B. cinerea*, in which the orthologous PEF-hand protein contributes to the virulence of environmental isolates of this phytopathogenic species (18).

Research on the membrane-protective role of PEF-hand proteins in the context of host-pathogen interactions has also been shown in human cells, which can be attacked by numerous microbial membrane-pore forming toxins or mechanical forces that disrupt host cell membranes (19–24). In their notable study, Westman *et al.* demonstrated that the human PEF-hand protein, ALG-2, maintains epithelial cell integrity during membrane attack by invasive hyphae of *C. albicans* that secrete the pore-forming peptide toxin, candidalysin (23). ALG-2 is a pre-formed Ca^2+^ sensor protein that mediates a membrane repair response by initiating assembly of the ESCRT protein complex, which seals membrane wounds by removing the damaged membrane area (25–27). In human and fungal cells, the membrane recruitment of PEF-hand proteins is dependent on external Ca^2+^ (16, 26), indicating that the increase in cytosolic Ca^2+^ levels after membrane wounding acts as a common localized signal to activate these membrane stress responses. However, other similarities and possible differences in the membrane repair mechanisms used by different types of eukaryotic cells, including diverse fungi, remain unknown.

In this study, we investigated the role of PEF-hand proteins in promoting fungal membrane integrity and virulence using *C. albicans* as a representative of human- pathogenic fungi (8). As part of the human microbiome, *C. albicans* can cause opportunistic infectious disease, which remains challenging to combat given the morphological plasticity and metabolic adaptability of this species, and the occurrence of resistance to antifungal drugs (28–30). During the commensal and pathogenic lifestyles of this polymorphic fungus, yeast and hyphal cells can encounter membrane stress caused by antifungal immunity (12, 31). In particular, the constitutively-polarized invasive hyphae of *C. albicans* require that membrane integrity is tightly controlled at their ergosterol-rich cell tips, which expand by incorporating new membrane lipids and proteins through vesicle exocytosis (32, 33). In the model organism, *Saccharomyces cerevisiae,* the Pef1 ortholog plays a role in the polarized growth phase of dividing yeast cells (34). Here, we investigated Pef1 function during polarized growth of *C. albicans* yeast cells and filaments, in the presence and absence of ergosterol-perturbing drugs. The study revealed that Pef1 localizes to sites of membrane perturbation, which includes sites of membrane re-organization during polarized growth and sites of membrane-damage caused by applied compounds. Therefore, the function of Pef1 in *C. albicans* is consistent with the role of PEF-hand proteins in eukaryotic cell membrane-repair, but is of additional significance in this pathogen, where the plasma-membrane is seen as a fungus-specific target for novel antifungal-drug development.

## Results

### Pef1 is required for polarized growth and resistance to SDS and Ca^2+^-depletion in yeast cells of *C. albicans*

Using PEF1 from *N. crassa* as a query for a BLASTp search, we identified an uncharacterized PEF-hand protein in *C. albicans*, which we hence designate Pef1. As annotated in the *Candida* Genome Database (http://www.candidagenome.org/), the diploid genome of *C. albicans* contains two identical alleles of a 1116-bp intronless gene (C2_08020C_A and C2_08020C_B), which encode the Pef1 protein with a length of 371 amino acids (aa). A BLASTp search with the full-length aa sequence of Pef1 from *C. albicans* as a query revealed that this predicted PEF-hand protein shares approximately 38 % and 29 % of total identity with the *N. crassa* PEF1 and *S. cerevisiae* Pef1 proteins, respectively (34, 16). Moreover, Pef1 in *C. albicans* is ∼ 33 % identical to the human cytosolic Ca^2+^ sensor protein, ALG-2, which initiates plasma membrane repair during mechanical and chemical forms of membrane disruption (23, 26). Pef1 also shows ∼ 34 % identity with peflin, another human PEF-hand protein that interacts with ALG-2 in the context of ER-to-Golgi transport (35). Functional predictions and aa alignments of these PEF-hand proteins indicated that the five helix-loop-helix motifs, which form the Ca^2+^-binding PEF-hand region, are conserved between the human and fungal orthologs (Fig. S1A). In contrast to ALG-2, however, peflin and the fungal orthologs contain an additional N-terminal extension with predicted disordered regions that share a relatively low percentage of aa sequence identity or similarity in between each other as compared to their conserved PEF-hand regions (Fig. S1B).

To functionally characterize Pef1 in *C. albicans*, we generated a set of control and mutant strains (Fig. S2), which included a mutant lacking the *PEF1* gene (*pef1*Δ) and a fluorescently tagged reporter strain that expressed a *GFP-PEF1* construct in the *pef1*Δ mutant (*pef1*Δ + *GFP-PEF1*). When grown in YPD broth, *pef1*Δ achieved a lower final OD_600_ value at 30 or 37 °C than the wild-type *PEF1* control strain, SN78-R (Fig. 1A-B). Although initially delayed in growth, the complemented *pef1*Δ + *GFP-PEF1* strain reached levels of growth that were similar to those of SN78-R after ∼ 3 h (Fig. 1A-B), suggesting that two functional copies of *PEF1* are required for normal growth. Microscopy revealed that the *pef1*Δ mutant showed a cell separation defect during yeast cell budding, which was also reversed by expression of *GFP-PEF1* in the complemented strain, demonstrating functionality of the tagged copy of Pef1 (Fig. 1C). Consistent with a proposed role of Pef1 in polarized growth in *S. cerevisiae* (34), the GFP-Pef1 fusion protein in *C. albicans* was also found to localize at the bud tip or the bud neck of the dividing yeast cells grown in broth (Fig. 1D). Since establishing and maintaining cell polarity are Ca^2+^-dependent processes (36), we next tested whether loss of the PEF-hand protein affects growth under Ca^2+^-limited conditions. Compared to the control strain, deletion of *PEF1* resulted in reduced growth on YPD agar plates containing the Ca^2+^ chelator, EGTA, which was rescued by supplementation of the media with Ca^2+^, or by complementation with *GFP-PEF1* (Fig. 1E). Although the yeast cell separation defect had previously been interpreted as a defect in cell wall integrity (34), the *pef1*Δ mutant was only slightly more susceptible to the cell wall-disrupting agent, calcofluor white (CFW), than the control strains. In contrast, exposure to sodium dodecyl sulfate (SDS), a surfactant that primarily disrupts cell membrane integrity, strongly impaired growth of the *pef1*Δ mutant (Fig. 1F). Since SDS also impacts cell wall integrity (37), we supplemented the medium with sorbitol, which osmotically stabilizes cells with cell wall defects (38). However, sorbitol did not reverse the specific growth defect of the *pef1*Δ mutant in the presence of SDS (Fig. S3). Taken together, these results indicate that Pef1 in *C. albicans* is involved in the maintenance of cell membrane integrity and functions at sites of polarized growth in yeast cells.

**Fig. 1:**
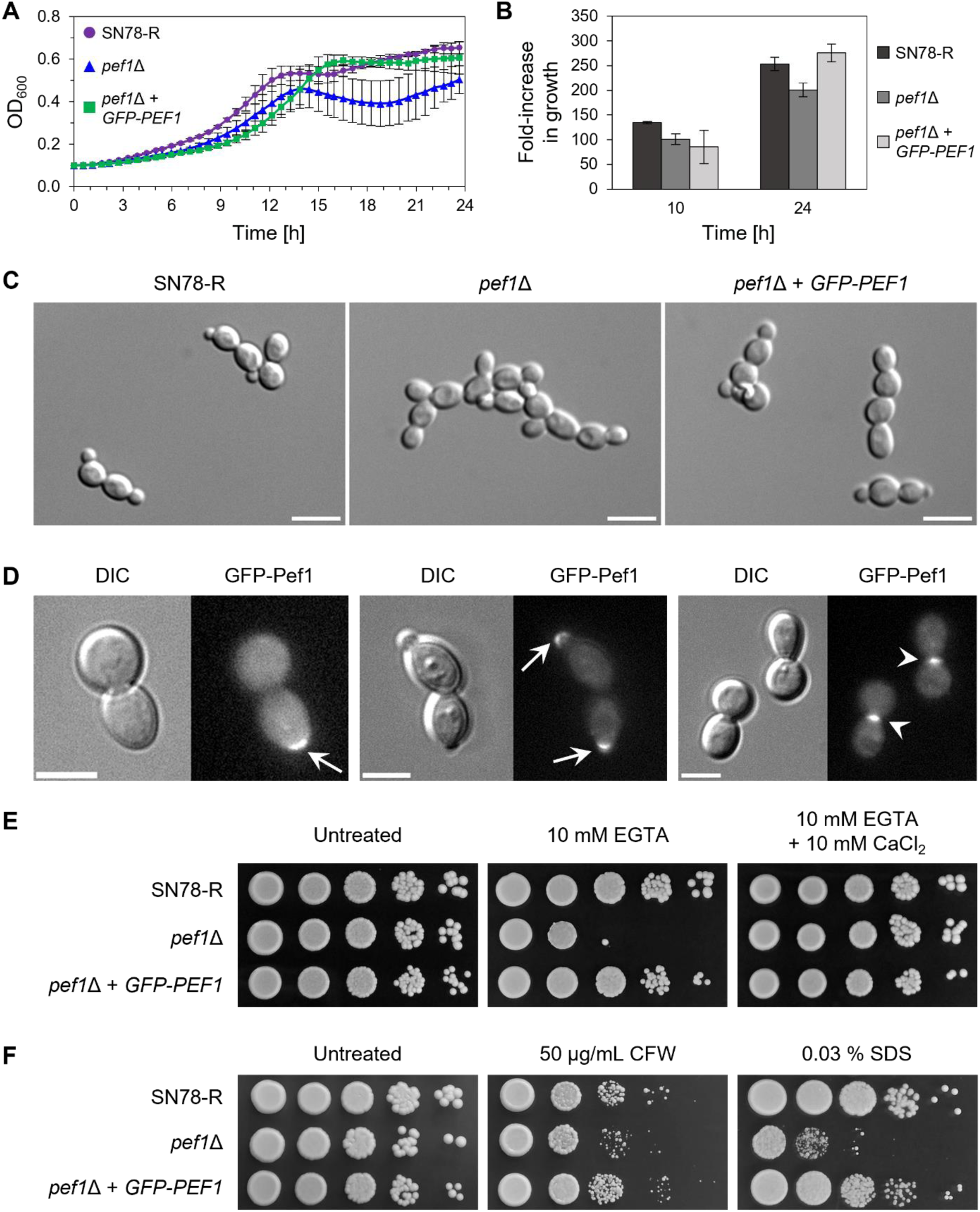
Pef1 supports polarized growth and resistance to SDS and EGTA in *C. albicans* yeast cells. **A:** Growth of the control strain SN78-R (MW-Ca81), the *pef1*Δ mutant (MW-Ca27) and the *pef1*Δ strain complemented with a *GFP-PEF1* construct (MW-Ca58) for 24 h at 30°C in YPD broth in microtiter plates. The growth curves show the mean OD_600_ values with errors (standard deviation, Std Dev) represented as bars from two biological replicates. **B:** Measurement of growth of the strains from panel A at 37°C in YPD broth in shaking flasks. The OD_600_ values were normalized for each strain and the relative fold-increase in growth was determined at the indicated time points. The bars represent the mean values with errors (Std Dev) from two biological replicates. **C:** Differential interference contrast (DIC) images of budding yeast cells from the strains described in panel A after 3 h of incubation at 30°C in YPD broth. Scale bars: 10 μm. **D:** DIC and fluorescence microscopy images of yeast cells from the GFP-Pef1 reporter strain (MW-Ca58) in synthetic defined (SD) medium supplemented with 5 mM Ca^2+^. GFP-Pef1 is localizing at the cortex of the emerging daughter cells (arrows) and at the neck in between the dividing cells (arrowheads) during yeast cell budding. Scale bars: 5 μm. **E:** Colonies of the strains indicated in panel A grown in the presence or absence of the Ca^2+^-chelator EGTA. Ten-fold serial dilutions of yeast cells were spotted onto YPD agar plates with the indicated supplements and incubated for 2 d at 37°C. **F:** Growth of colonies as described in panel E on YPD agar plates with and without calcofluor white (CFW) or sodium dodecyl sulphate (SDS). Images of the colonies were captured after 2 d of incubation at 37°C.

### GFP-Pef1 localizes at the hyphal tip and is required for plasma membrane integrity in extending apical compartments

We next examined the localization of GFP-Pef1 during the filamentous growth of *C. albicans*. In serum, GFP-Pef1 localized to hyphal tips and also to septa (Fig. 2A), consistent with a function for Pef1 at sites of polarized growth and membrane remodeling. Deletion of *PEF1* did not affect the emergence or extension rate of hyphal filaments (Fig. S4A-B). However, the mutant hyphae showed signs of stress and damage as indicated by aberrant vacuole morphology and distribution, in addition to cell bursting near the hyphal apex (Fig. S4B-C). Staining of filaments with propidium iodide (PI), a red-fluorescent, DNA-binding dye that only enters cells with a compromised plasma membrane (39), indicated that the apical hyphal compartments of the *pef1*Δ hyphae were permeable to this compound (Fig. 2B). Approximately one third of the mutant filaments were PI-positive (Fig. 2C), and this primarily occurred in the apical compartments, with subapical compartments remaining PI-negative (Fig. 2D). In contrast, the complemented mutant formed hyphae that were morphologically indistinguishable from the control strain and exhibited negative PI-staining (Fig. 2B-C). Treatment of *pef1*Δ hyphae with the lipophilic, red-fluorescent dye FM4-64 showed that ∼ 31 % (58/188) of *pef1*Δ hyphae internalized FM4-64 within 15 min compared to only ∼ 2 % (2/96) of the hyphae of the SN78-R control strain, in which the dye predominantly labelled the cell periphery at this time point (Fig. S4C). Overall, these observations suggest that the polarized localization of Pef1 is required to maintain integrity of the plasma membrane during normal hyphal growth of *C. albicans*.

**Fig. 2:**
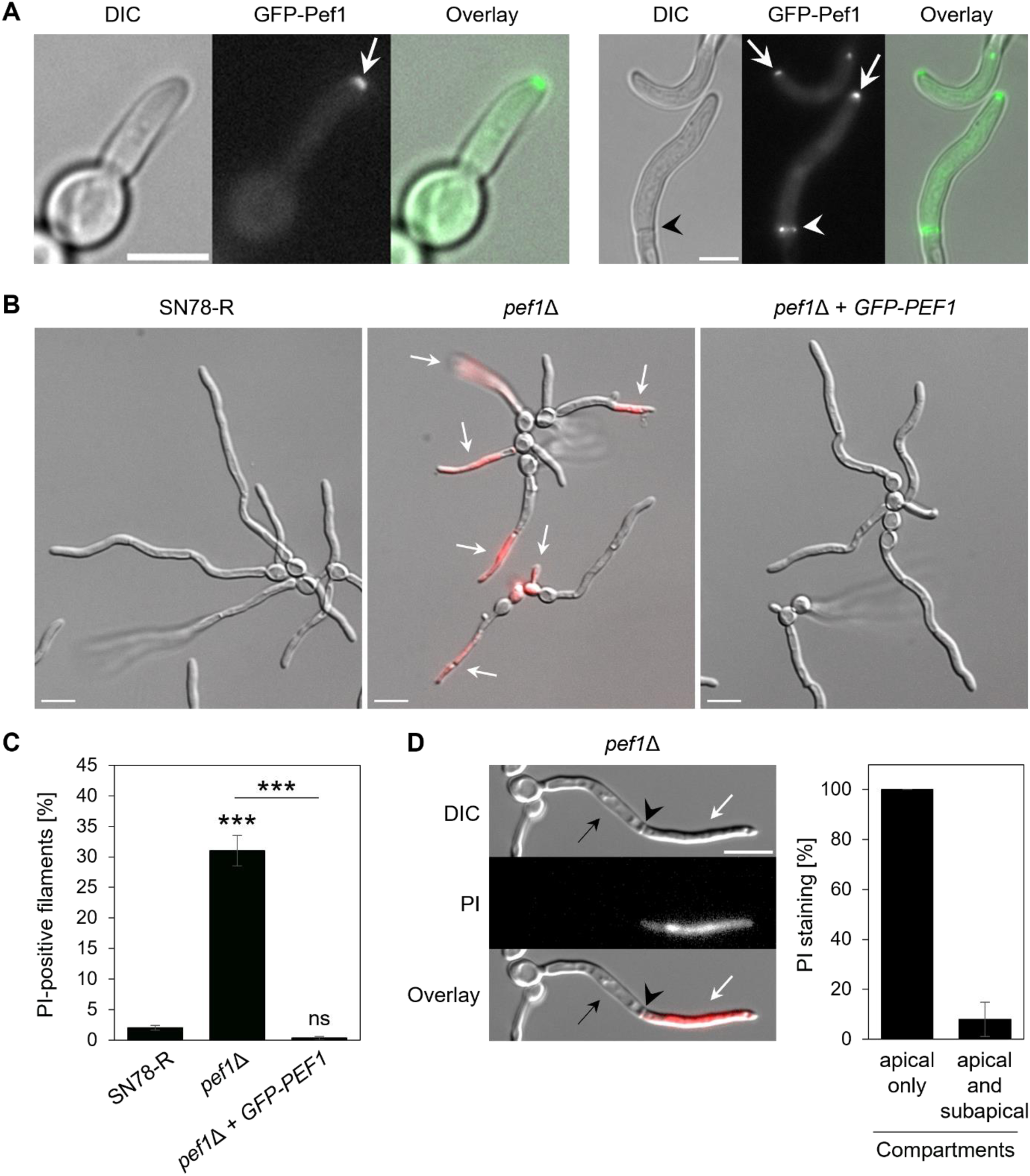
Pef1 localizes at the hyphal tip and facilitates membrane integrity of *C. albicans* hyphae. **A:** Incubation of the GFP-Pef1 reporter strain (MW-Ca58) in 20% fetal bovine serum (FBS) to induce the formation of filaments. Images were captured by fluorescence microscopy after 1 h (left panel) and 3 h (right panel) of incubation at 37°C. GFP-Pef1 is localizing at the hyphal tip (arrows) and at a septum (black and white arrowheads). Scale bars: 5 μm. **B:** Overlay images (DIC and fluorescence) of filaments formed by SN78-R (MW-Ca81), the *pef1*Δ mutant (MW-Ca27) and the complemented strain (MW-Ca58) after 3 h of incubation at 37°C in FBS. Staining with propidium iodide (PI) reveals accumulation of the red-fluorescent dye in the filaments of the *pef1*Δ mutant (arrows). Scale bars: 10 μm. **C:** Quantification of the percentage of filaments from the strains show in panel B that stained positive with PI after 3 h of incubation at 37°C in FBS. The black bars show the mean values with errors (Std Dev) from three independent experiments per strain. Statistically significant differences (***, p < 0.001; ns, not significant) were assessed by one-way ANOVA analysis with Tukey’s correction for multiple comparisons. **D:** Quantification of PI staining patterns in septated PI-positive filaments of the *pef1*Δ mutant after 3-4 h of incubation at 37°C in FBS. Left: DIC and fluorescent images of a representative PI-positive *pef1*Δ filament with one septum (arrowhead) separating an apical (white arrow) and subapical (back arrow) compartment. Scale bar: 10 μm. Right: Percentage of PI staining in apical and/or neighboring subapical hyphal compartments, with mean values and error bars (Std Dev) retrieved from a total of 52 septated hyphae in 4 independent experiments.

The ability of *C. albicans* to form filaments in liquid media can be separated from its capacity to form hyphae that invade semisolid substrates (40). Although hyphae of the *pef1*Δ mutant showed a defect in sustaining apical integrity in liquid serum medium (Fig. 2B), the filaments formed by colonies did not show an altered ability to penetrate solid serum agar and were able to reach agar invasion depths similar to the control strains when incubated at 30°C or 37°C (Fig. S5A-B). However, on solid Spider medium there was a delay in the macroscopic appearance of filamentation in the *pef1*Δ mutant at 30 °C, but not at 37 °C (Fig. S5C). Nevertheless, Spider agar invasion at 30°C or 37°C was not affected by the deletion of *PEF1* (Fig. S5D-E). Furthermore, incubation in liquid Spider medium did not reveal any significant increase in PI staining of the filaments of the mutant in contrast to the defect it showed in serum (Fig. S6A-B). These results suggest that Pef1 is required for promoting membrane integrity during hyphal growth in serum, as opposed to non-physiologically-relevant media.

### The impaired hyphal integrity of the calcineurin mutant correlates with an altered tip localization of Pef1

It is well-known that mutants lacking a functional Ca^2+^/calcineurin-Crz1 signaling pathway are impaired in cell integrity during normal growth and are hypersensitive to membrane-damaging compounds (41–43). Consistent with previous studies on the importance of the catalytic subunit of the calcineurin phosphatase, Cna1, for *C. albicans* survival in serum (41), hyphae of the *cna1*Δ mutant stained largely positive with PI when grown in serum, with levels of staining exceeding those observed for *pef1*Δ (Fig. 3A-B). In contrast, the filaments of the *crz1*Δ mutant, which lacks the transcription factor downstream of calcineurin (43), were indistinguishable from those of the control strains, SC5314 and CAI-4/CIp10 (Fig. 3A-B). These observations indicate that the processes mediating membrane integrity during normal filamentous growth of *C. albicans* in serum are dependent on Pef1 and calcineurin, but independent from Crz1-mediated gene expression.

**Fig. 3:**
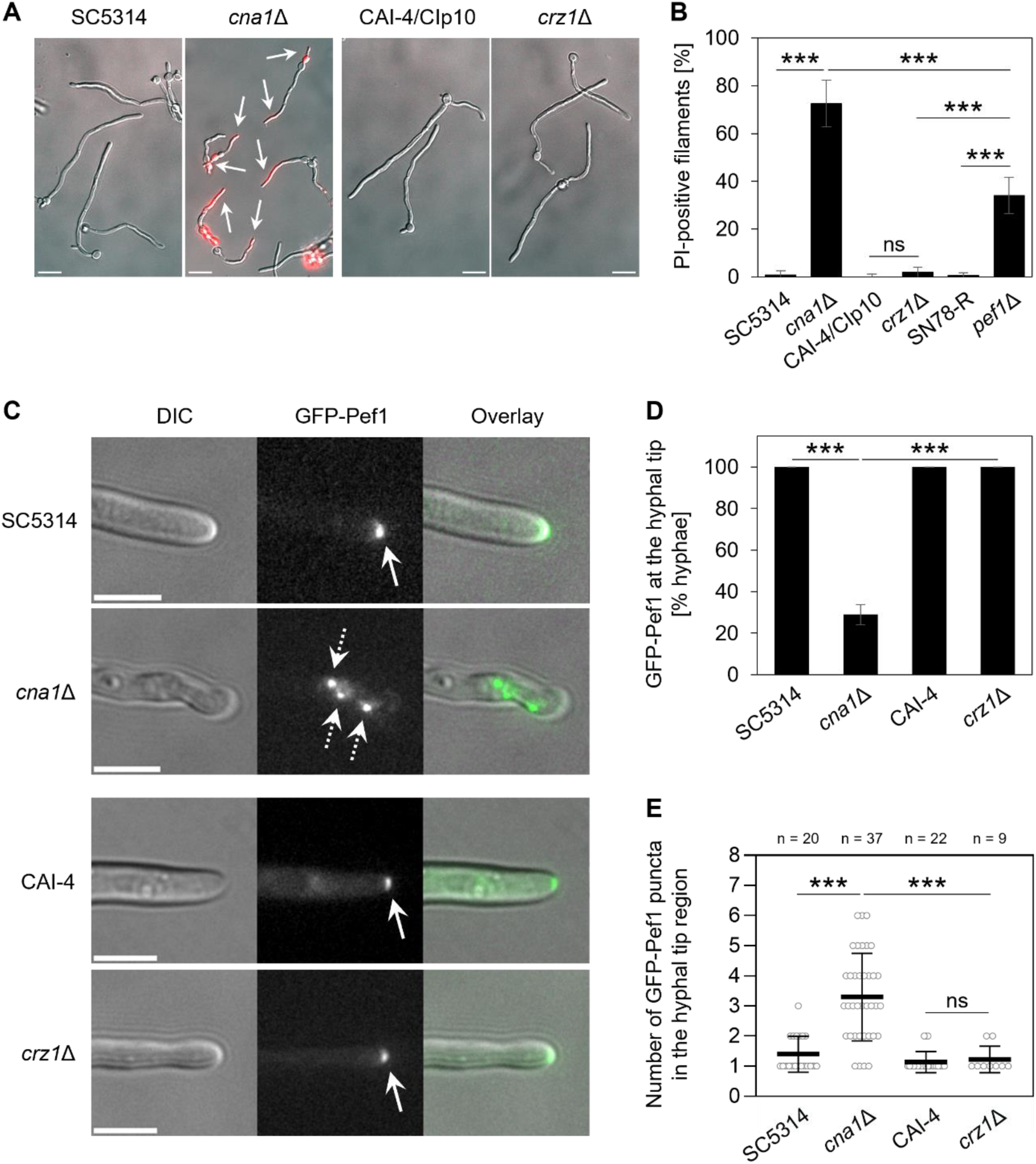
Loss of calcineurin impairs hyphal integrity and alters the localization pattern of Pef1. **A:** Overlay images (DIC and red fluorescence) of hyphae of the *cna1*Δ (SCCMP1M4) and *crz1*Δ (MKY380) mutants and their respective reference strains (SC5314 and CAI-4/CIp10). Filamentation was induced by incubating yeast cells for 3 h at 37°C in 20 % FBS followed by staining with PI. Arrows indicate PI-positive hyphae. Scale bars: 10 μm. **B:** Comparison of the percentage of filaments showing PI staining after growth in FBS in mutants lacking the *CNA1*, *CRZ1* or *PEF1* gene next to their respective control strains. The black bars show mean values with errors (Std Dev) from three to seven technical replicates derived from two independent experiments per strain. **C:** Localization of GFP-Pef1 in hyphae of the *cna1*Δ (MW-Ca134) and *crz1*Δ (MW-Ca132) mutants and their respective control strains SC5314 and CAI-4 expressing the fluorescent fusion protein (MW-Ca124 and MW-Ca125). After incubation for 3 h at 37°C in 20 % FBS, the protein is focused at the hyphal tips of both control strains and the *crz1*Δ mutant (solid arrows), whereas several punctate signals appear in a wider region at the hyphal tip in the *cna1*Δ mutant (dotted arrows). Scale bars: 5 μm. **D:** Percentage of filaments from the strains presented in panel C showing a focused localization of GFP-Pef1 at the hyphal tip. The black bars show the mean values with errors (Std Dev) from two to three technical replicates per strain. **E:** Quantitative analysis of the number of GFP-Pef1 puncta in the hyphal tip region of the strains shown in panel C. The scatter plot represents the mean values and error bars (Std Dev) of the number of fluorescent signals scored in each hyphal tip region (open circles) among the total number (n) of filaments for each strain. Statistically significant differences between the mean values shown in B, D and E (***, p < 0.001; ns, not significant) were assessed by one-way ANOVA analysis with Tukey’s correction for multiple comparisons.

To identify a functional link between calcineurin and Pef1, we expressed GFP-Pef1 during hyphal growth of the *cna1*Δ and *crz1*Δ mutants. In hyphae of the controls, SC5314 and CAI-4, localization of GFP-Pef1 in the tip region was the same as that previously observed (Fig. 2A & Fig. 3C). GFP-Pef1 also localized to the hyphal tip in both the *cna1*Δ and *crz1*Δ mutants but the fluorescent signal at the apex was strongly reduced only in the calcineurin-deficient strain (Fig. 3C-D). Strikingly, GFP-Pef1 also appeared as dispersed puncta in the hyphal tip area of *cna1*Δ (Fig. 3C & E). Thus, the contrasting hyphal integrity phenotypes of the *cna1*Δ and *crz1*Δ mutants correlated with differential localization patterns of GFP-Pef1. These observations suggest that calcineurin activation could be partially responsible for the localization of Pef1 during hyphal growth or that loss of calcineurin allows the development of membrane damage that requires localized Pef1 activity for repair.

### Pef1 puncta are redistributed within the plasma-membrane on exposure to membrane-damaging compounds

To explore the requirement for localized Pef1 membrane repair activity further, we exposed *C. albicans* hyphae to the membrane-interacting compounds, amphotericin B (AmB, a polyene), or tomatine (a saponin). In both cases, the intensity of the GFP-Pef1 signal at the hyphal tip was reduced and the signal fragmented into puncta within the apical zone (Fig. 4A-C). Moreover, additional puncta of GFP-Pef1 appeared in subapical regions. Overall, these observations suggest that Pef1 localizes to the hyphal tip during polarized growth but is redistributed to points throughout the plasma-membrane during chemically-induced whole-cell membrane disruption.

**Fig. 4:**
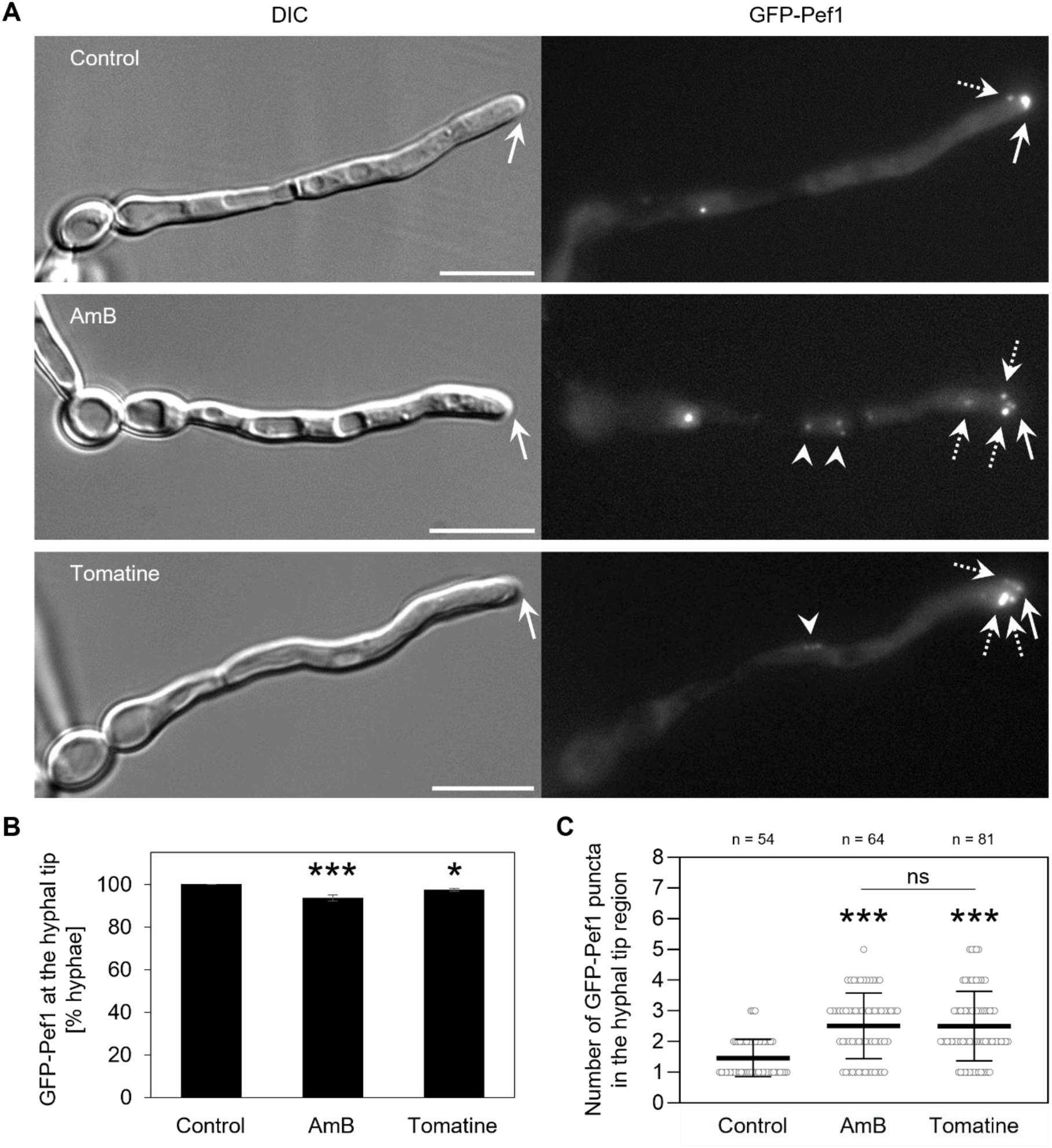
Membrane-disrupting antifungal compounds alter the hyphal tip localization of Pef1. **A:** Localization of GFP-Pef1 (reporter strain MW-Ca58) in hyphae during membrane stress caused by antifungal compounds. The filaments were grown for 3 h at 37°C in 20 % FBS prior to treatment with 2 μg/mL amphotericin B (AmB) or 50 μg/mL tomatine. Images were captured after about 5 to 15 min of antifungal exposure. Compared to the focused signal of GFP-Pef1 at the center of the hyphal tip (solid arrow) in the control filament, several additional puncta of GFP-Pef1 appear at the hyphal tip region (dotted arrows), and some also in regions distal to the hyphal tip (arrowheads). Scale bars: 10 μm. **B:** Quantification of the percentage of filaments as shown in panel A with a focused localization of GFP-Pef1 at the hyphal tip. In the control and during the treatment with AmB or tomatine, almost all filaments show a fluorescent signal of GFP-Pef1 at the polarized cell tips. The black bars represent the mean values with errors (Std Dev) from two to four independent experiments per strain. **C:** Quantitative analysis of the number of distinct signals (puncta) of GFP-Pef1 in the hyphal tip region in the absence and presence of AmB or tomatine. From the filaments analyzed in panel B, the number of puncta scored for each hyphal tip (open circles) is represented in a dot plot, including mean values and error bars (Std Dev) from the total number (n) of hyphae analyzed per condition. Statistically significant differences in B and C (***, p < 0.001; *, p < 0.05; ns, not significant) were assessed by one-way ANOVA analysis with Tukey’s correction for multiple comparisons.

As the localization pattern of Pef1 to the hyphal tip during polarized growth is very similar to that seen for polarisome components, we next tested whether the polarisome marker, Spa2, is redistributed in response to chemical insult in the same manner (44). Both proteins showed a similar polarized localization in untreated hyphae but, unlike GFP- Pef1, the Spa2-GFP distribution was unaffected by the antifungal compounds, albeit with a slightly reduced signal in AmB-treated hyphae (Fig. S7A-C). These observations suggest that, while both Spa2 and Pef1 localize to the hyphal apex to during polarized growth, this is achieved via different mechanisms and Pef1 is unlikely to be a constitutive member of the polarisome complex.

### Pef1 function is required for resistance to membrane-damaging compounds

We next examined the distribution of GFP-Pef1 in yeast cells during treatment with AmB or tomatine. Similar to our observations in hyphae, exposure to the polyene and saponin induced the formation of fluorescent puncta throughout the peripheral membrane of yeast cells (Fig. 5A). Thus, both types of ergosterol-binding compounds dramatically altered the localization pattern of Pef1 in yeasts and hyphae, possibly due to recruitment of the protein to sites of chemically-induced membrane disruption.

**Fig. 5:**
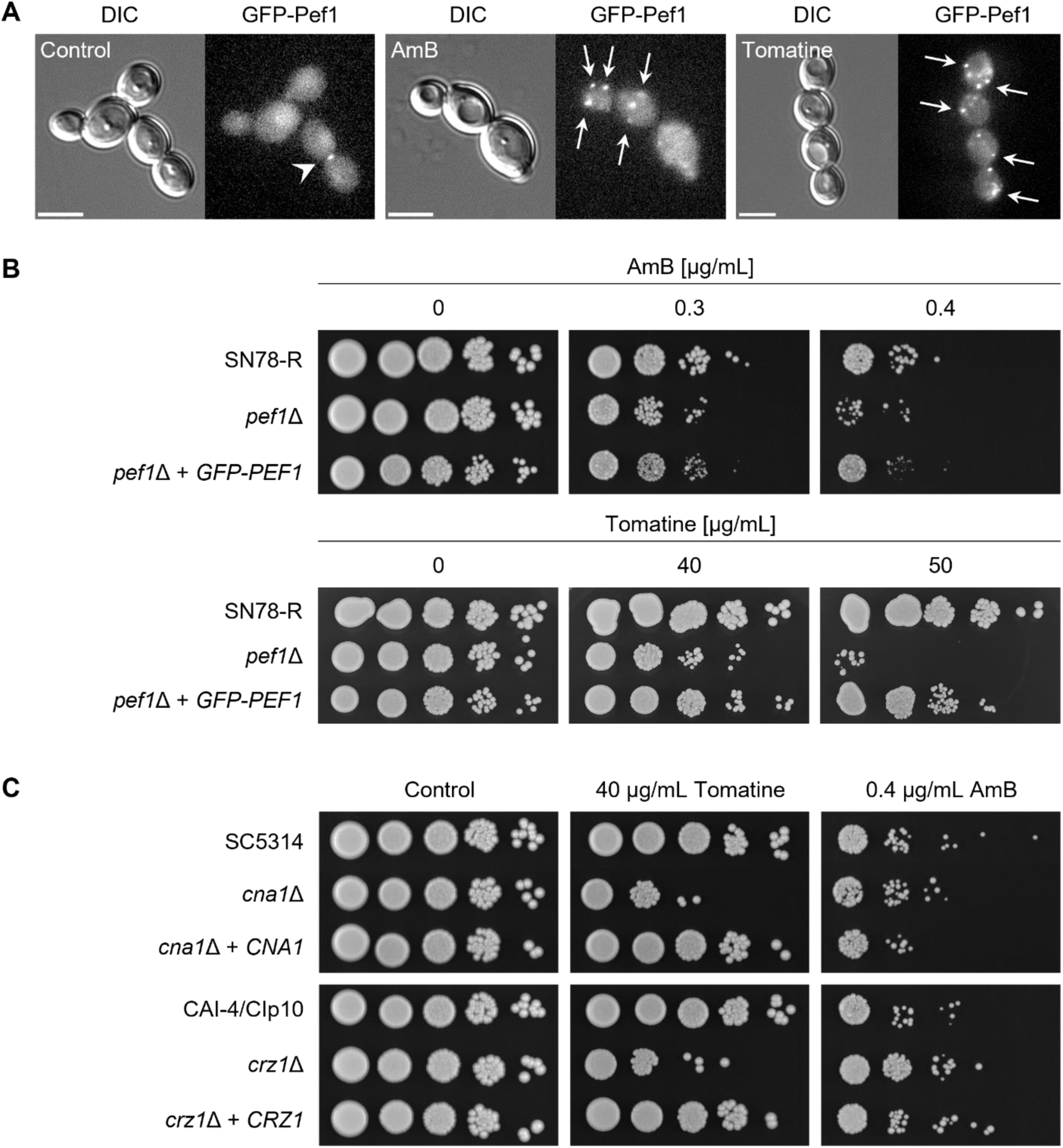
Pef1 and Ca^2+^/calcineurin signaling contribute in a similar manner to the adaptation to membrane-disrupting fungicides. **A:** Localization of GFP-Pef1 (reporter strain MW-Ca58) during the treatment of yeast cells with antifungal compounds. Yeast cells from YPD cultures were washed in SD medium supplemented with 5 mM Ca^2+^ prior to the exposure to 2 μg/mL AmB, 50 μg/mL tomatine or 0.125 % DMSO (control). Images were captured after 5 to 15 min of antifungal treatment. While GFP-Pef1 localizes at sites of cell polarity in the control (arrowhead; see also Fig. 1D), the protein accumulates at the cell periphery and at intracellular sites during antifungal treatment (arrows). Scale bars: 5 μm. **B:** Colonies grown from 10-fold serial dilutions of yeast cells of the control strain SN78-R (MW-Ca81), the *pef1*Δ mutant (MW-Ca27) and the complemented strain (MW-Ca58) spotted on solid YPD medium with and without the indicated concentrations of AmB or tomatine. Growth was captured after 2 d of incubation at 37°C. **C:** Ten-fold serial dilutions of yeast cells from the *cna1*Δ (SCCMP1M4) and *crz1*Δ (MKY380) mutants as well as the respective control strains (SC5314, CAI-4/CIp10) and complemented mutants (SCCMP1MK2, MKY381) were spotted onto YPD agar plates with and without tomatine or AmB and incubated for 2 d at 37°C.

We also evaluated the importance of Pef1 function in countering their antifungal effect and compared this with the requirement for calcineurin and the Crz1 transcription factor. Spot assays showed that the *pef1*Δ mutant was only mildly susceptible to AmB compared to the control strains (Fig. 5B). In a similar manner, there was no change in susceptibility to the polyene-derivative, nystatin, even though this commonly-used topical antifungal also induced the formation of multiple puncta of GFP-Pef1 in yeast cells (Fig. S8A-B). In contrast, the *pef1*Δ mutant showed clear susceptibility to tomatine (Fig. 5B). Interestingly, the *pef1*Δ mutant shared these different fungicide susceptibility phenotypes with both mutants of the calcineurin signaling pathway (Fig. 5C). Taken together, these results demonstrate a differential role of Pef1 in *C. albicans* in response to antifungal compounds that cause direct membrane disruption that is comparable to that of PEF1 in *N. crassa* (16), indicating that orthologous PEF-hand proteins in distantly related fungal species mediate a conserved response to membrane damage.

### Pef1 is required for virulence in the Galleria mellonella infection model

To determine whether Pef1 function is required for fungal pathogenicity, we tested the virulence of the *pef1*Δ mutant in the larvae of the greater wax moth, *Galleria mellonella*, a validated model for fungal virulence studies (45). This model features innate immune responses that include the release of membrane-targeting antimicrobial peptides (46, 47). One week after infecting groups of larvae with *C. albicans*, the *pef1*Δ mutant produced a population death rate of ∼ 50 %, compared to 100 % and ∼ 80 % of the larvae infected with SN78-R and the complemented *pef1*Δ+GFP-PEF1 strain, respectively. Statistical analysis confirmed that the percentage of survival of larvae infected with the *pef1*Δ mutant was significantly higher compared to those infected with the control strains (Fig. 6). For comparison, when infecting *G. mellonella* with the *cna1*Δ and *crz1*Δ mutants, we observed death rates of ∼ 20 % and ∼ 40 %, respectively, compared to ∼ 50 – 90 % in the corresponding control strains (Fig. S9A-B), which is in agreement with the impaired virulence reported for these mutants in murine models of systemic candidiasis (41, 43, 48, 49). Loss of Pef1 therefore attenuates the pathogenicity of *C. albicans* in this infection model to similar levels seen for mutants defective in the calcineurin signaling pathway. This is consistent with the susceptibility and Pef1 localization data across these mutants, supporting the view that Pef1 plays an important role in maintaining membrane integrity, both at polarity sites and during membrane stress imposed by external factors.

**Fig. 6:**
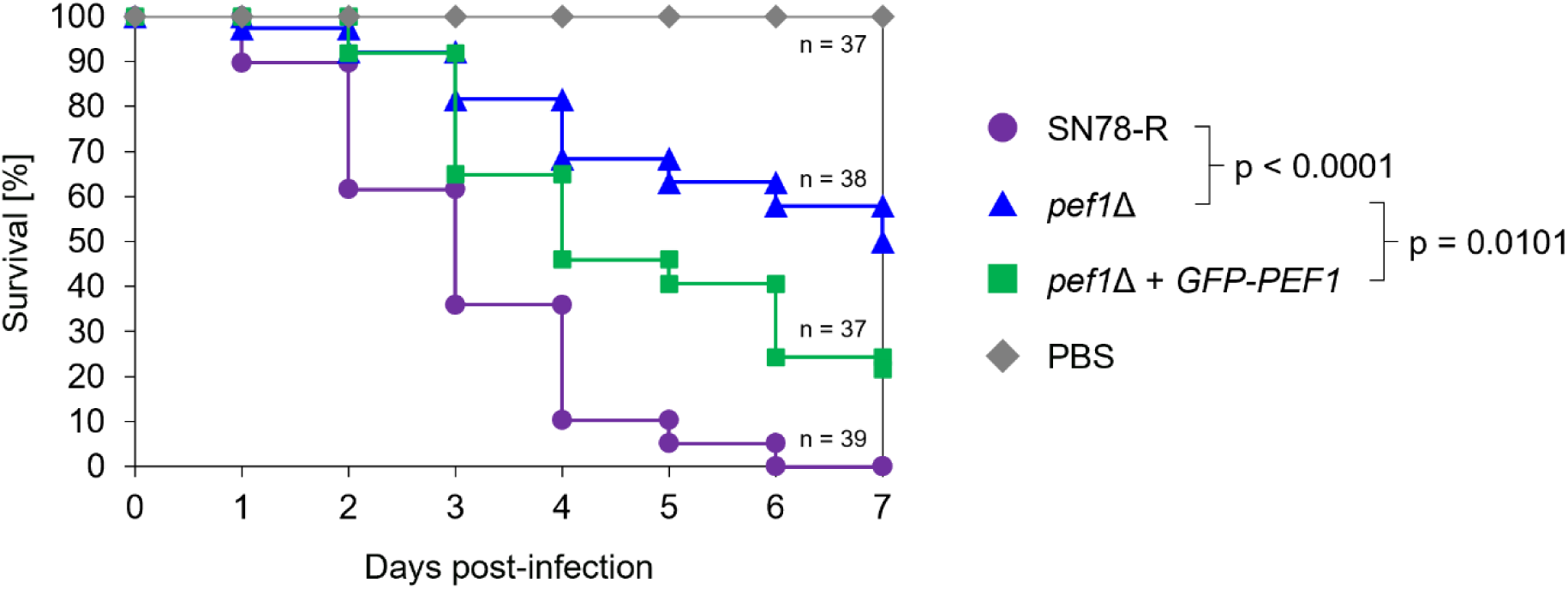
**Loss of Pef1 attenuates the virulence of *C. albicans.*** Survival plots of *G. mellonella* larvae infected with yeast cells of SN78-R (MW-Ca81), the *pef1*Δ mutant (MW-Ca27) or the complemented strain (MW-Ca58). After injection with 1x10^6^ yeast cells (prepared in phosphate buffered saline, PBS) or with sterile PBS as a control, the larvae were incubated for 7 days at 37°C and checked daily for survival. The plots represent pooled data from four independent experiments with the indicated total number (n) of larvae used per strain. Comparisons of survival plots were statistically assessed by Log-rank analysis resulting in the indicated p values.

## Discussion

Fungal PEF-hand proteins have previously been associated with polarized growth during yeast cell budding in *S. cerevisiae* and cell membrane repair in germlings and filaments of *N. crassa* and *B. cinerea* (16, 18, 34). In this study, we demonstrate that the orthologous Pef1 protein in *C. albicans* shares both functions through supporting polarized growth and maintaining overall membrane integrity in the yeast and hyphal forms of this polymorphic species. Our study contributes to the growing evidence on the highly conserved role of eukaryotic PEF-hand proteins in response to membrane disruption, which in fungi is currently best understood in the context of membrane-targeting antifungal compounds that trigger the fast and dynamic recruitment of these Ca^2+^-binding proteins to the cell membrane. In addition to this conserved function in maintaining membrane integrity, Pef1 in *C. albicans* takes an additional role during polarized growth, which seems to be specific for fungal species with a yeast morphotype. Overall, our results suggest that the *C. albicans* Pef1 protein is recruited to cell sites that are undergoing membrane perturbation, whether this is caused by normal membrane re- organization during polarized growth or abnormal damage by external factors. Pef1 appears to act as a specialist ‘first responder’ that stabilizes the plasma membrane at sites of perturbation, where immediate repair is required to maintain cell viability. Its function is therefore relevant to normal fungal growth, survival of immune-system attack and resistance mechanisms against membrane-targeting antifungal drugs.

### The *C. albicans* PEF-hand protein Pef1 is required for membrane integrity during hyphal growth

In contrast to filamentous fungi, commonly known as molds, in which PEF-hand proteins reside in the cytosol during normal hyphal growth (16, 18), the *C. albicans* Pef1 protein localized at the polarized hyphal tip when growing in the filamentous morphotype. In yeast cells, Pef1 was similarly recruited to sites of cell polarity during budding, consistent with previously reported localization patterns for the orthologous protein in *S. cerevisiae* (34). This role of Pef1 in yeast-forming fungal species is evidenced by the cell separation defect that mutants lacking Pef1 have in common in both species, suggesting that the membrane is compromised, or at least disorganized, at septa, such that the molecular organization of abscission cannot occur. Strikingly, one of the highlights in our study is the finding that hyphae of the *C. albicans pef1*Δ mutant were specifically impaired in maintaining membrane integrity during polarized growth in serum, but not in agar plates containing low serum concentrations or other lab-derived media. The lack of a hyphal integrity defect in liquid Spider medium suggests that the specific hypha-inducing conditions impact whether hyphal integrity is dependent on Pef1 or not. These observations indicate that the fungal plasma-membrane in serum-induced hyphae is more prone to damage during polarized growth than when cells are grown in non-physiological, synthetic media.

In this respect, our finding that apical cells in *pef1*Δ hyphae, but not in yeast or germinating hyphae, became non-viable in serum is similar to findings for the *cna1*Δ mutant. Consistent with previous research, calcineurin is required for the survival of *C. albicans* and other *Candida* species in serum, which represents a form of Ca^2+^ ion stress (41, 50–53). Interestingly, we found that only the *cna1*Δ, but not the *crz1*Δ, mutant was deficient in maintaining membrane integrity during hyphal growth in serum, and that these different phenotypes correlated with different localization patterns of Pef1 at the hyphal tip. These observations indicate important roles of non-canonical calcineurin signaling and Pef1 function during normal hyphal growth in serum. Although the *cna1*Δ mutant shared this defect with the *pef1*Δ mutant, the much larger degree of impaired hyphal integrity by loss of Cna1 function likely reflects the broad role of Ca^2+^/calcineurin signaling in promoting cellular integrity on both the plasma membrane and the cell wall levels (15). Since the *pef1*Δ mutant was barely impaired by cell wall stress, we conclude that non-canonical calcineurin signaling and Pef1 have functionally overlapping roles in maintaining membrane integrity during normal growth.

### Fungal PEF-hand proteins share a conserved function in maintaining cell membrane integrity during membrane disruption by antifungal compounds

All so-far studied PEF-hand proteins in fungi share a conserved role in the response and adaptation to external conditions that cause acute membrane disruption. Our results in *C. albicans* support pervious findings in *N. crassa* and *B. cinerea* that showed the quick membrane recruitment of those Pef1 orthologs after treatment with a polyene or saponin (16, 18). Although Pef1 is constitutively associated with polarized membrane sites in yeast and hyphal cells of *C. albicans*, the dramatic change in its subcellular localization upon exposure to membrane-disrupting fungicides underscores its primary role in mediating a membrane stress response. As discussed above, the localization patterns of Pef1 in untreated dividing yeast cells and at the growing hyphal tip might be a specific function in cell polarity that could be related to a continuous response to intrinsic forms of membrane stress during normal growth. We propose that the altered localization of Pef1 during antifungal-induced stress represents a sudden shift from guarding membrane integrity during polarized growth towards initiating a repair response after membrane attack.

The dynamic recruitment of fungal PEF-hand proteins reflects the intensity and severity of membrane attack by fungicides that directly interact with ergosterol (3, 7). The plasma membrane of polarized growing hyphal tips in different fungal species is highly enriched in ergosterol (18, 33, 54–56), indicating that PEF-hand proteins localize to membrane sites where maximum damage occurs. Moreover, it is remarkable that mutants lacking fungal PEF-hand proteins in different species share the same differential phenotypes in the presence of polyenes and saponins. Similar to the filamentous fungi, the *pef1*Δ mutant of *C. albicans* did not show increased susceptibility to nystatin or AmB despite the membrane recruitment of Pef1 in response to these polyenes. In contrast, Pef1 was required for the tolerance of tomatine, a saponin compound that this human microbiome-associated fungus unlikely encounters in the host environment. Such similar observations in distantly related fungal species promote the idea that the molecular basis of Pef1- mediated membrane stress adaptation in fungi is highly conserved despite their different ecologies and lifestyles. It is possible that *C. albicans* encounters other metabolites or molecules present in its niches, either originating from host cells or microbes, that have membrane-disrupting modes of action similar to those of saponins, which could explain the conservation of this response in this species.

Many membrane-targeting compounds likely have in common that they cause the formation of smaller or bigger membrane pores or ruptures, which ensue a sudden influx of external Ca^2+^ into the cytosol. It is well known that the quick recruitment of the human ALG-2 protein to membrane wounds after mechanical or chemical forms of plasma membrane disruption is dependent on an increase in cytosolic Ca^2+^ (26, 27). In *N. crassa*, the availability of external Ca^2+^ is also required for the membrane recruitment of PEF1 (16), implying that this basic function of PEF-hand proteins is likely conserved in all eukaryotic cells. Given the strong conservation of the Ca^2+^-binding helix-loop-helix motifs in human and fungal PEF-hand proteins, we assume that fungal orthologs act in a similar manner as Ca^2+^-sensor proteins as previously established for ALG-2. Interestingly, it was recently shown that Ca^2+^ directly stimulates the membrane binding of ALG-2 by neutralizing electrostatic repulsion with acidic phospholipid membranes (57). This Ca^2+^ binding mechanism might also mediate the membrane association of the fungal PEF-hand proteins, given the strong conservation of the EF-hand motifs. Moreover, the disordered N-termini of fungal ALG-2 orthologs might also allow for a direct or indirect membrane association, since these regions are known to enable close membrane associations and the interaction with other proteins that might be directly tethered to the lipid bilayer (58). ALG-2 does not feature such disordered N-terminal extension, but its interaction partner, peflin, does (35, 59). While it is unknown whether peflin is directly associated with membrane repair, we hypothesize that important differences in the mobilization, interactions and functions of fungal and human PEF-hand proteins exist.

### Pef1-mediated membrane integrity is a specific subset of Ca^2+^-dependent mechanisms that support cellular integrity

In addition to the membrane recruitment of PEF-hand proteins, a rapid rise in cytosolic Ca^2+^ levels also activates Ca^2+^/calcineurin signaling, which mediates a broad adaptive membrane stress response in fungi (15). Since Ca^2+^ influx is a common signal, we reasoned that Pef1 function is, at least in part, related to this major stress response pathway. Our results showed that the *pef1*Δ mutant shares many fungicide-related phenotypes with mutants defective in Ca^2+^/calcineurin signaling. Loss of either Cna1 or Crz1 caused hypersensitivity to tomatine, but not to AmB, which largely resembled the phenotypes of the *pef1*Δ mutant. This implies that canonical Ca^2+^/calcineurin signaling and Pef1 function are individually dispensable for survival during the exposure to polyenes, but each process is required for the tolerance of saponin-induced membrane disruption. We assume that other forms of direct membrane disruption also induce both mechanisms, which jointly operate to reach the full scale in the response to membrane attack.

Controlling Ca^2+^ homeostasis is crucial for establishing and maintaining cell polarity, and Ca^2+^ uptake at the hyphal tip controls the growth direction during tip extension (60–62). In addition to passive Ca^2+^ influx during sporadic membrane disruption at the growing hyphal tip region, the active and controlled form of Ca^2+^ uptake might also activate both calcineurin and Pef1 in a simultaneous or sequential manner. In any case, both molecular responses might be mechanistically linked when membrane integrity is diminished. Our observation that punctate spots of GFP-tagged Pef1 localized at the hyphal tip region of the *cna1*Δ mutant likely indicates the recruitment of this protein to sites of membrane damage during growth rather than a mislocalization of Pef1 in the absence of calcineurin signaling. Since this altered localization pattern partially resembled the puncta induced by fungicides in a calcineurin-competent strain, we conclude that these subcellular dynamics of Pef1 further highlight its primary role in the membrane stress response.

Both the PEF-hand protein and calcineurin are likely part of a larger network of functionally redundant and independent cellular mechanisms that are mounted during passive or active forms of Ca^2+^ influx. We assume that Pef1 covers a specific subset of those functions in maintaining membrane integrity, which Ca^2+^/calcineurin signaling mediates to a larger extent. While Ca^2+^ ions are key for the activation of both processes, it is possible that the specific origin, form, magnitude and/or duration of Ca^2+^ influx determines which specific membrane integrity mechanisms are induced. Exploring cellular Ca^2+^ dynamics during membrane stress with real-time reporter systems, such as the genetically-encoded Ca^2+^ indicator GCaMP6 (63, 64), would improve our mechanistic understanding on how Ca^2+^-dependent processes specifically mediate the adaption to growth conditions and antifungal compounds that challenge membrane integrity. Moreover, the expression of fluorescently tagged Pef1 as a reporter in additional sets of mutant strains as well as clinical isolates of *C. albicans* that show different membrane stress phenotypes would complement studies on Ca^2+^ dynamics and collectively expand our knowledge on Ca^2+^-dependent mechanisms of membrane integrity.

### The functions of Pef1 are highly relevant for the virulence of *C. albicans*

Based on the above-discussed roles of Pef1 in polarized and membrane integrity, we reasoned that these functions, either alone or in combination, are highly relevant for the pathogenicity of *C. albicans*. The filamentous growth of this human pathogen is a major determinant of its pathogenic potential, given that invasive hyphae disrupt epithelial cell barriers and the hypha-enforced lytic escape from the phagolysosome kills macrophages (29, 65, 66). However, mutants of *C. albicans* with defects in polarized growth *in vitro* do not necessarily show attenuated virulence in infection models (40). Our data indicate that Pef1 supports membrane integrity during normal hyphal growth, but that it is dispensable for filamentation and agar invasion. Since the host environment contains factors that can directly disrupt microbial membranes, in particular membrane-targeting antimicrobial peptides that are part of innate immunity and conserved in *G. mellonella* larvae (13, 45), we tested in this invertebrate infection model the virulence of the *pef1*Δ mutant. Our observation that loss of Pef1 significantly attenuated the killing of infected insect larvae identifies this PEF-hand protein as a factor contributing to the pathogenic potential of *C. albicans*. Compared to mutants defective in Ca^2+^/calcineurin signaling, this pathogenicity defect of the *pef1*Δ mutant appears less pronounced. This likely reflects the much broader implications of Ca^2+^/calcineurin signaling in promoting cellular stress adaptation (15), whereas Pef1 covers a much more specific role in response to direct membrane stress. Nonetheless, the relative contribution of Pef1 to pathogenicity in this model is remarkable and implies that maintaining fungal membrane integrity by this protein and yet-to-identify associated molecular factors and mechanisms is highly relevant in the context of host- pathogen interactions. Just as Pef1 of *C. albicans* likely responds to membrane attack elicited by the immune system, it was previously demonstrated that human cells employ an ALG-2-initiated membrane repair process when attacked by candidalysin (23). This indicates that during fungal infection, a protective response towards the reciprocal usage of membrane attack involves defense mechanisms mediated by PEF-hand proteins with conserved functions in membrane repair. During the complex and dynamic interactions of these combating eukaryotic cells, quickly adapting to direct and acute forms of membrane damage is vital for either side. The dynamic mobilization of cytosolic factors, including the Ca^2+^ influx-triggered membrane recruitment of PEF-hand proteins, to immediately protect the barrier function of the plasma membrane would ensure cellular survival as slower subsequent mechanisms of promoting and restoring membrane integrity are still being mounted and employed. Strong evidence supporting a model for the concerted activation of pre-formed and induced molecular factors that protect the cell membrane has recently been presented in *B. cinerea* during its interaction with tomato plants (18). While the membrane-targeting host defense compound tomatine triggers as a first-level response the quick recruitment of the BcPEF1 ortholog to the fungal membrane, the induction of several membrane-modifying and tomatine-detoxifying enzymes required transcriptional activation. However, the availability of these second- level mechanisms to protect the fungal membrane strongly promoted tomatine tolerance and plant pathogenicity in a mutant strain lacking BcPEF1 (18). Thus, progressively unfolding different mechanisms of membrane integrity can boost the pathogenic potential of fungi. It remains to be investigated which molecular mechanisms other than the Pef1- mediated response are involved in the adaptation of *C. albicans* to membrane stress.

Both conserved and unique molecular factors are expected to contribute to membrane repair and integrity in the context of host-pathogen interactions. Moreover, these mechanisms might also be relevant for the adaption of *C. albicans* and other human-pathogenic fungi to antifungal drugs used in the treatment of patients with fungal disease. Considering the increasing incidence of resistance to antifungal drugs (67), the molecular dissection of membrane integrity mechanisms could inform novel strategies to combat hard-to-treat fungal infections. Further research on membrane repair in fungi could identify currently unknown points of vulnerability on the membrane level of pathogenic species, which might be exploited to design novel and specific treatment options against fungal infections, and to prevent and combat antifungal resistance.

## Materials and Methods

### Standard growth conditions, media and reagents

Yeast cell suspensions of *C. albicans* were grown as standard overnight cultures at 30°C and 200 rpm in YPD broth (1 % [w/v] yeast extract [BD Difco], 2 % [w/v] bacto-peptone [BD Difco], 2 % [w/v] glucose), unless otherwise stated. Cell concentrations were photometrically determined as optical density at 600 nm (OD_600_). Prototrophic and auxotrophic strains were selected on synthetic defined (SD) medium (0.67 % [w/v] yeast nitrogen base [YNB; BD Difco], 2 % [w/v] glucose, 1.5 % [w/v] agar) with or without the supplementation of histidine, leucine and/or uridine (each at 200 µg/mL), respectively. To induce hypha formation, yeast cells were washed in sterile MilliQ (MQ) water or double-distilled water and incubated at 37°C in 20 % (v/v) fetal bovine serum (FBS; Sigma) using 8-well polymer chamber slides with ibiTreat surface modification (ibidi). Stock solutions of antifungal compounds and stress agents were prepared as follows: 10 % (w/v) SDS in MQ water; 10 mg/mL CFW (fluorescence brightener 28, Sigma) in MQ water; 1 to 2 mg/mL AmB (Sigma) in DMSO; 50 mg/mL nystatin (Millipore) in DMSO; 20 or 25 mg/mL tomatine (biorbyt or Carl Roth) in either DMSO (for microscopy) or methanol with 0.2 % (v/v) formic acid (for plate assays). For molecular cloning of plasmids, electro-competent or chemo-competent *Escherichia coli* strains were grown at 37°C in Luria-Bertani medium (1 % [w/v] tryptone [BD Difco], 0.5 % [w/v] yeast extract [BD Difco], 0.5 % [w/v] NaCl) supplemented with 100 µg/mL ampicillin or 50 µg/mL kanamycin. Standard protocols were used for plasmid DNA manipulation, bacterial transformations and the preparation of plasmid DNA from bacterial cell suspensions or genomic DNA from fungal yeast cell suspensions.

### C. albicans strain construction

All *C. albicans* strains used and constructed in this study are listed in Tab. 1. Oligonucleotides (primers) used for molecular cloning and confirmatory PCR analyses are listed in Tab. 2.

**Tab. 1:**
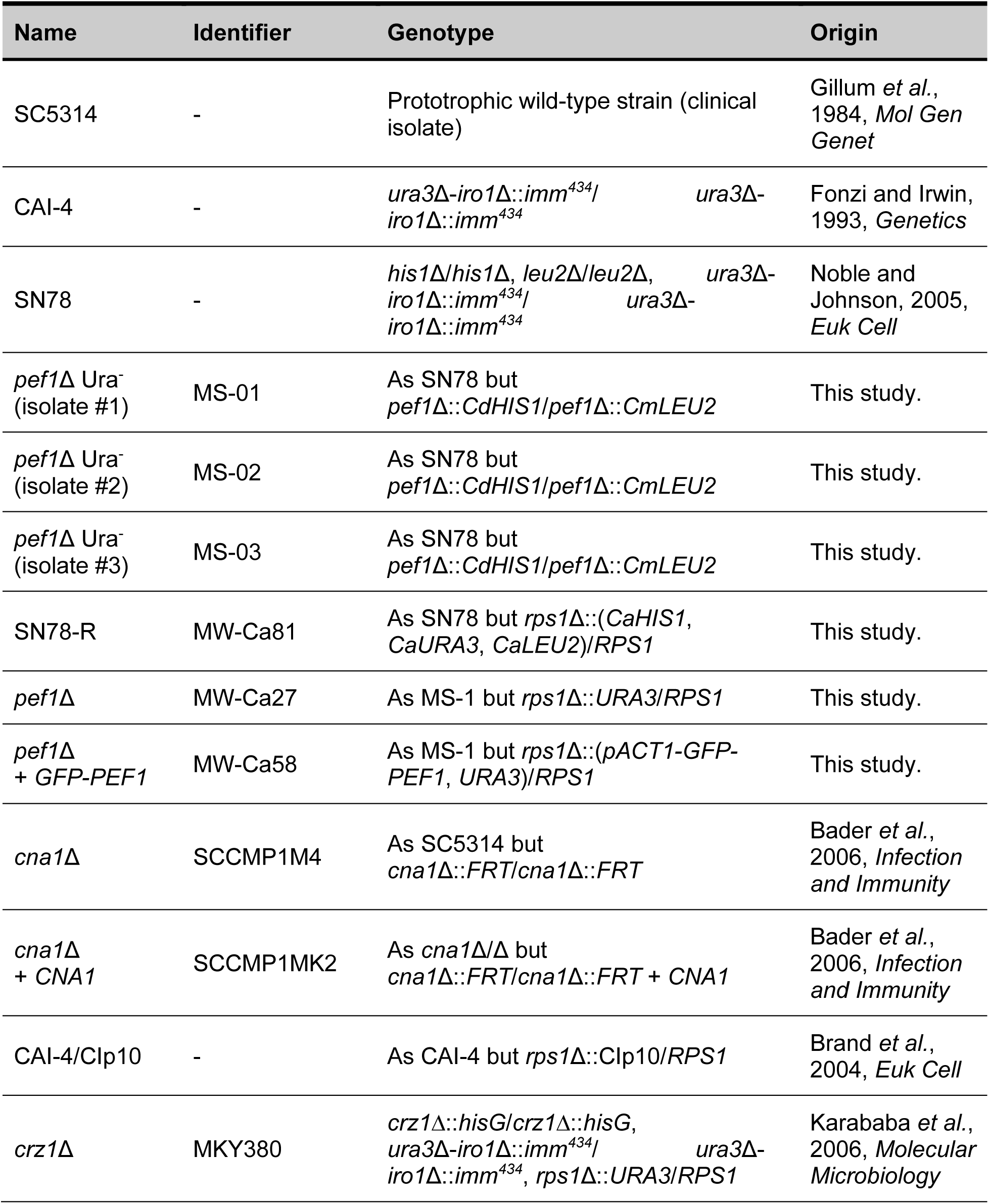
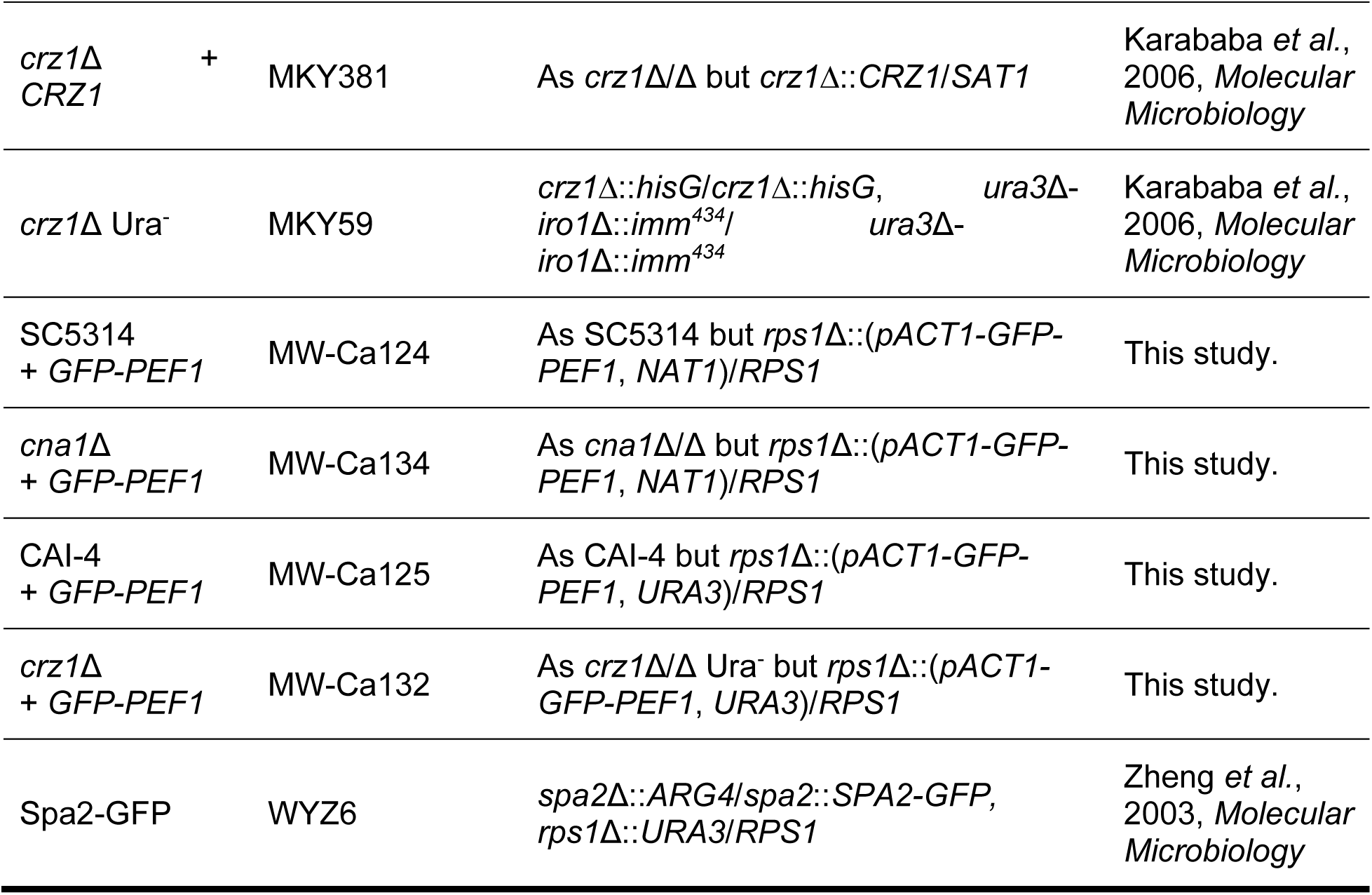
Strains of *C. albicans* used in this study.

**Tab. 2:**
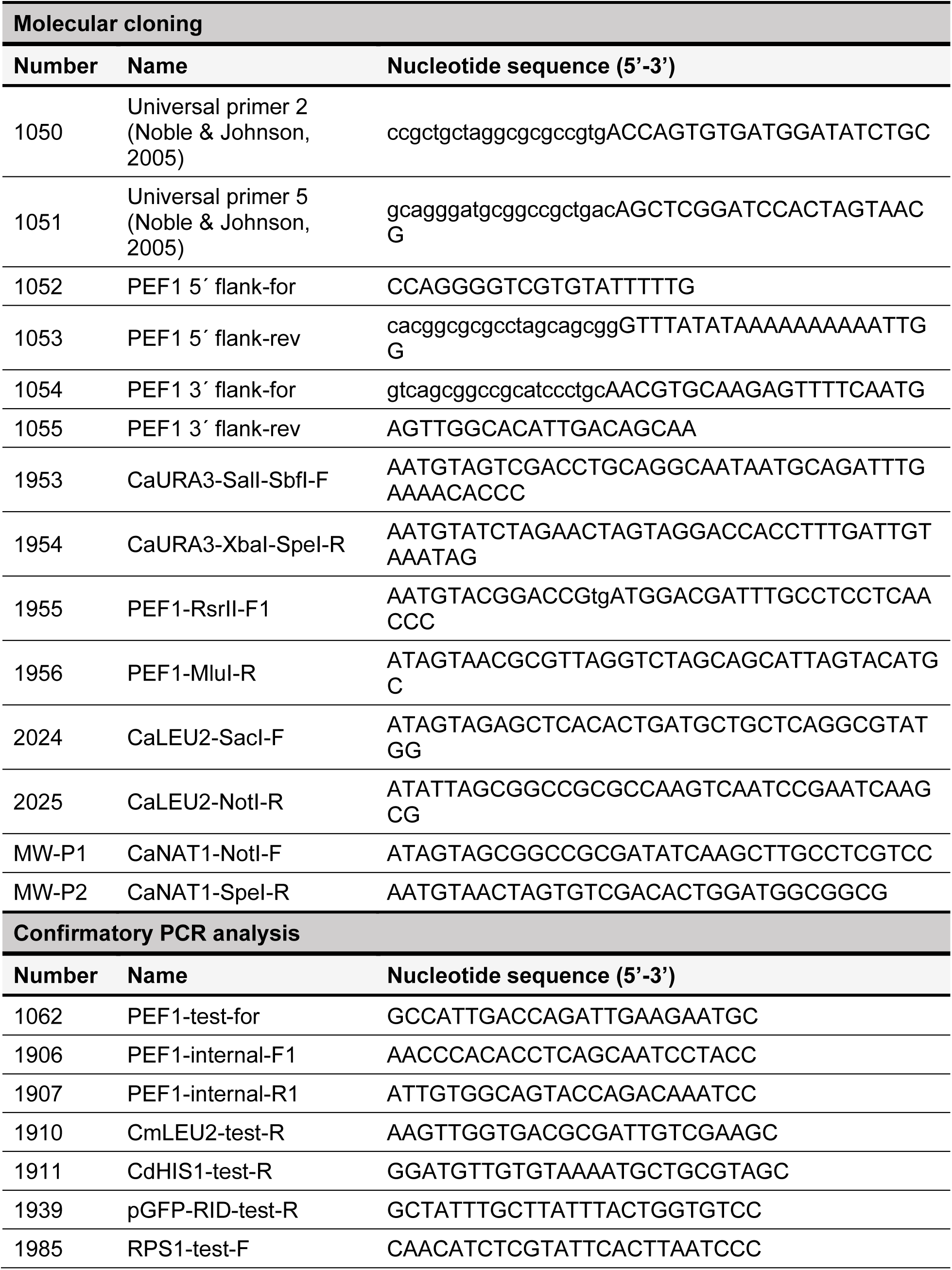
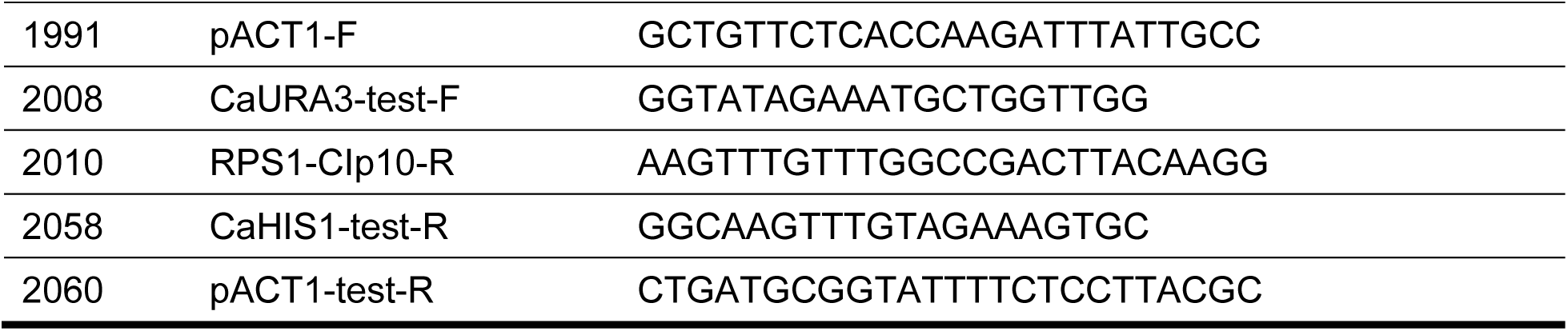
List of primers used in this study.

### Deletion of the PEF1 gene

To delete both alleles of the *PEF1* gene in the triple-auxotrophic (His*^-^*, Leu*^-^*, Ura*^-^*) reference strain, SN78, we employed a previously published gene deletion strategy (68). Using genomic DNA from strain SC5314 as a template, left (5’) and right (3’) flanks homologous to ∼ 350-bp sequences upstream and downstream of the *PEF1* gene were PCR-amplified with the primer pairs 1052/1053 and 1054/1055, respectively. The selectable markers *LEU2* (from *Candida maltose*, *CmLEU2*) and *HIS1* (from *Candida dubliniensis*, *CdHIS1*) were PCR-amplified from plasmids pSN40 and pSN52, respectively, with the primer pair 1050/1051 as described before (68). To generate both *PEF1* knock-out (KO) cassettes, the 5’ and 3’ flanks were combined with the *LEU2* or *HIS1* marker using a fusion PCR approach (68). Yeast cells of strain SN78 were prepared for transformation with the KO cassettes using a lithium acetate/DTT protocol and transformed by electroporation as described before (69, 70). First, the *LEU2*- containing KO cassette was transformed into SN78 followed by selection of heterozygous mutants on SD agar plates supplemented with uridine and lacking leucine. Second, one of these heterozygous isolates was further transformed with the *HIS1*-containing KO cassette, resulting in homozygous *pef1*Δ Ura^-^ mutants (MS-01 to MS-03) that were selected on uridine-containing SD agar plates lacking both leucine and histidine (Fig. S2A-B). Strain MS-01 was transformed with the StuI-linearized plasmid CIp10 to integrate a copy of *URA3* at the *RPS1* locus as describe before (71, 72), resulting in the prototrophic *pef1*Δ mutant strain, MW-Ca27 (Fig. S2C). To restore prototrophy in the triple-auxotrophic parental strain SN78, plasmid CIp20, which contains the markers *URA3* and *HIS1* from *C. albicans* (73), was linearized with SacI and NotI to allow for the integration of the *LEU2* marker from *C. albicans* (*CaLEU2*, including ∼ 850 bp of its promoter and ∼ 200 bp of its terminator regions in analogy to *CmLEU2* from pSN40). The *CaLEU2* marker was PCR-amplified from SC5314 with the primer pair 2024/2025. In line with the CIp series of vectors for integration at the *C. albicans RPS1* locus (71, 73), the resulting plasmid was named CIp40 (lab identifier 1122/B342; Fig. S2D). Strain SN78 was transformed with StuI-linearized CIp40, as described above, to generate the prototroph SN78-R (strain MW-Ca81). After selection on SD agar plates lacking leucine, histidine and uracil/uridine, integration of CIp40 at the *RPS1* locus was confirmed by PCR (Fig. S2D).

### GFP-tagging of Pef1

The *pef1*Δ Ura^-^ mutant (strain MS-01) was complemented with a construct that encodes Pef1 fused at its N-terminus to the *C. albicans*-codon optimized sequence of the green fluorescent protein (GFP) for live-cell imaging and fluorescence microscopy (74). Plasmid pExpArg-pACT1GFPRID (75) was stepwise modified for expression of the *GFP-PEF1* construct under the *ACT1* promoter (*ACT1*p) (76). First, the XbaI and SalI sites were used to replace the *ARG4* marker by the *URA3* gene from SC5314, which was PCR-amplified with the primer pair 1953/1954 (resulting in the intermediate plasmid pExpUra-pACT1GFPRID, lab identifier 1102). Next, the *RID* reporter was excised via RsrII and MluI and replaced by the *PEF1* gene including 500 bp of the sequence downstream of its stop codon, which was PCR-amplified from SC5314 with the primer pair 1955/1956. The resulting plasmid pExpUra-pACT1-GFP-PEF1 (lab identifier 1104/B341; Fig. S2E) and linearized with StuI prior to transformation as describe above. Transformants were selected on SD agar plates without uracil/uridine. *GFP-PEF1* integration at the RPS1 locus was confirmed by PCR (Fig. S2E) and fluorescence microcopy. Plasmid pExpUra-pACT1-GFP-PEF1 was integrated into the reference strain, CAI-4, and the *crz1*Δ Ura^-^ mutant, and confirmed by PCR. Integration of *GFP-PEF1* into SC5314 and the *cna1*Δ mutant (SCCMP1M4): the *URA3* marker in plasmid pExpUra- pACT1-GFP-PEF1 was excised by NotI and SpeI and replaced by a codon-optimized *NAT1* selectable marker (77). This *CaNAT1* marker was PCR-amplified using primer pair MW-P1/MW-P2 from plasmid CIp-NAT, a variant of plasmid CIp10 containing the *TEF1*p*- CaNAT1-TEF1*t construct (78). The resulting plasmid, pACT1-GFP-PEF1-NAT1 (lab identifier 1130/B343), was linearized with StuI and transformed into SC5314 and *cna1*Δ using a 15-min heat shock step at 44 °C followed by a recovery phase in YPD broth supplemented with uridine for 4 h at 30°C and 200 rpm. Transformants were selected on Sabouraud dextrose agar plates (Oxoid) supplemented with 200 to 400 µg/mL nourseothricin (Jena Bioscience). The *GFP-PEF1*-expressing SC5314 and *cna1*Δ strains were confirmed by PCR and fluorescence microscopy.

### Quantification of yeast cell growth in broth

#### Microtiter plates

Cells were inoculated in 5 mL YPD broth and incubated at 30 °C and 200 rpm overnight. Cells from 1 mL of each pre-culture were pelleted for 60 s at 3500 rpm and washed twice in 1 mL of fresh YPD broth. Cell suspensions were diluted to OD_600_ = 0.1, and 200 µL or sterile YPD broth were added to the wells of 96-well flat-bottomed plates (Sarstedt), with 3 – 6 technical replicates per sample. The plate was sealed with a Breathe-Easy sealing membrane (Diversified Biotech) and incubated for 24 h at 30 °C in an Infinite 200 PRO plate reader (TECAN) with constant orbital shaking (amplitude: 3 mm). The OD_600_ was measured at 4 positions per well at 30-min intervals (orbital shaking; duration: 8 s, amplitude: 3 mm). Average growth curves were obtained by normalizing the OD_600_ values to the starting inoculum.

#### Shaking flask cultures

5 mL of cells from overnight cultures were washed in fresh YPD and adjusted to OD_600_ = 10. Flasks (250-mL) containing 50 mL fresh YPD broth were inoculated to OD_600_ = 0.1. Flasks were incubated at 37 °C and 200 rpm, and growth determined by measuring OD_600_ at various timepoints.

#### DIC and fluorescence microscopy

Live-cell imaging was acquired on an inverted Zeiss AxioObserver Z1 microscope equipped with DIC and fluorescence setups using a Plan-Neofluor 40X/1.3 numerical aperture oil immersion objective (Carl Zeiss), with a 16-bit CoolSNAP H2 charge-coupled-device (CCD) camera (Photometrics) within an incubation chamber (PeCon GmbH) at 37 °C. Images were acquired with the ZEISS ZEN software and processed in the Fiji freeware (https://fiji.sc/).

#### Analysis of yeast cell budding

Cells from standard overnight cultures were diluted 1:10 in fresh YPD and further incubated at 30 °C and 200 rpm. At specific time points, samples were prepared on standard microscope slides and observed by DIC microscopy.

#### Localization of GFP-Pef1

For localization studies in the yeast morphotype, cells were first diluted 1:100 in fresh YPD broth, further incubated at 30 °C and 200 rpm for about 5 h, and diluted 1:10 in SD medium supplemented with 5 mM Ca^2+^. Prior to preparing samples for fluorescence microscopy, aliquots of the cells were treated with 2 µg/mL AmB, 20 µg/mL nystatin, 50 µg/mL tomatine or 0.2 to 0.5 % (v/v) DMSO. Images of the cells were captured within 5-15 min after treatment. For hyphal localization of GFP-Pef1, yeast cells were washed in ddH_2_O, resuspended in 20 % (v/v) FBS to OD_600_ = 0.01, and seeded into ibidi chambers at 300 µL/well. Hyphae were treated with AmB or tomatine at the above-mentioned final concentrations by adding 100 µL of the compounds or DMSO (4X concentrated in 20 % [v/v] FBS) to the wells. Images were captured by fluorescence microscopy within 15 min of antifungal exposure.

#### Staining with PI or FM4-64

Yeast cells were transferred to 20 % (v/v) FBS or Spider medium (79) and incubated for 3-4 h at 37 °C in ibidi chambers in 300 µL/well. For the staining with PI (stock solution of 1 mg/mL in MQ water), 100 µL of fresh FBS medium or Spider medium containing the dye (4X concentrated in 20 % [v/v] FBS) were added to a final concentration of 2 µg/mL. Cells were stained with FM4-64 (stock solution of 100 µg/mL in MQ water) by adding 100 µL of FBS medium supplemented with the dye (4X concentrated in 20 % [v/v] FBS) to a final concentration of 0.1 µg/mL. Images were captured by fluorescence microscopy at 5 to 15 min after the addition of each dye.

#### Antifungal compound susceptibility testing

The OD_600_ values were measured for overnight cultures in YPD and cell concentrations adjusted to OD_600_ = 1 in sterile ddH_2_O water. Drops of 5 µL from 10-fold serial dilutions in ddH_2_O were spotted onto solid media with and without AmB (0.1-1.0 µg/mL), nystatin (2-5 µg/mL), tomatine (20-80 µg/mL), SDS (0.01-0.04%), CFW (20-80 µg/mL) or EGTA (5- 20 mM). Plates were incubated at 37 °C for 1-2 d and images captured.

### Agar invasion assays

Agar invasion was assessed as previously described (80). Yeast cells from overnight cultures were diluted to 2.5 x 10^4^ cell per 5-µl spot, and inoculate Spider agar or 10 % (v/v) serum (FBS) agar (1.5 % [w/v]) plates. Colony diameters were measured after incubation at 30 °C or 37 °C for 5-7 d. To determine the depth of agar invasion, cross-sections of colonies were prepared with a razor blade and imaged against a ruler. For each strain and growth condition, the ratio between the average colony diameters and average penetration depths was calculated.

### Insect larvae infection model

*C. albicans* strains were grown in 5 mL YPD broth for ∼ 22 h at 30 °C and 200 rpm. Cells were diluted 1:100 in fresh YPD and incubated as before. Cells were harvested at 3,000 rpm for 1 min and washed three times in PBS buffer prior to determining the cell numbers with a photometer and a hemocytometer. Groups of 10 or 12 similarly sized larvae of *G. mellonella* were infected via the left last pro-leg with 20 µl of PBS buffer containing 2 x 10^6^ yeast cells or 20 µL of sterile PBS with sterile single-use insulin syringes (30G x 1/2’’; B.Braun, Germany). Larvae were kept for 7 d at 37 °C in the dark and survival monitored daily. Larvae displaying extensive dark-brown pigmentation and loss of motility were scored as dead.

### Statistical data analysis

Statistical analysis was performed using GraphPad Prism version 9 for Windows (GraphPad Software, San Diego, CA, USA).

## Acknowledgements

The authors would like to thank the fungal research community for sharing *C. albicans* strains, plasmids and protocols: Suzanne Noble, Aaron Mitchell, Robert Arkowitz, Martine Bassilana, Antonio Serrano, Sascha Brunke, Bernhard Hube and Steven Bates. Darren Thomson for assisted with microscopy and Cameron Bedford gave technical assistance. We also appreciate the support from Dieter Jahn, Davina Hiller, Stefan Barthels and Alejandro Arce Rodriguez from the Institute of Microbiology at TU Braunschweig.

## Author contributions

M.W., M.R.S., A.C.B., and A.F. designed research; M.W., M.R.S., and U.B. performed research; M.W., M.R.S., A.C.B., and A.F. analyzed data; and M.W., A.C.B., and A.F. wrote the paper.

## Supplemental Figures

**Fig S1:**
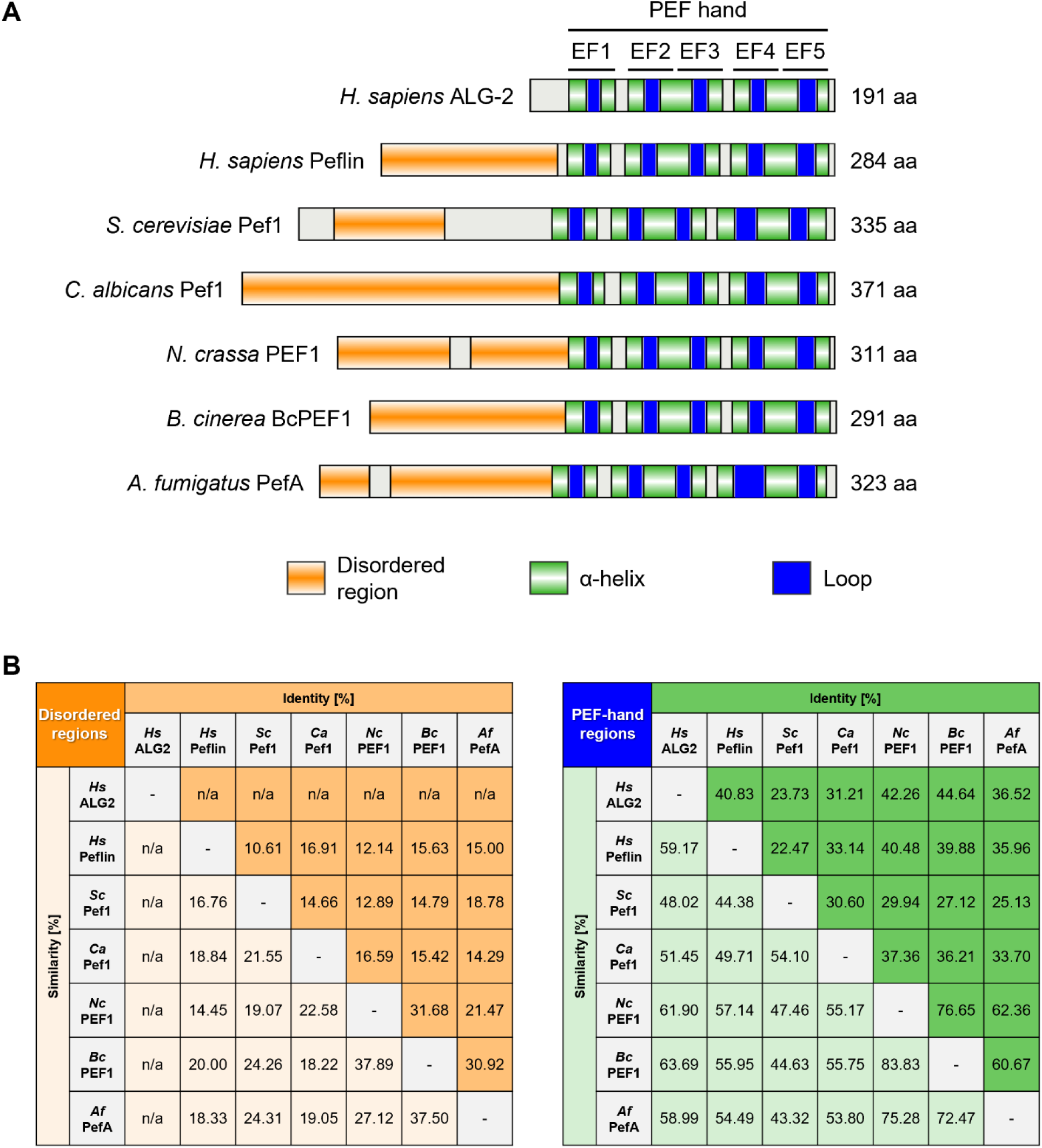
Alignment of mammalian and fungal PEF hand-proteins. **A:** Schematic representation of functional regions in the amino acid (aa) sequences of the *Homo sapiens* (*Hs*) proteins, ALG-2 and peflin (UniProt [https://www.uniprot.org/] entries: O75340 and Q9UBV8), and the orthologous penta-EF (PEF)-hand proteins from selected fungal species (FungiDB [https://fungidb.org/fungidb/app] entries: *Saccharomyces cerevisiae* [*Sc*], YGR058W; *Candida albicans* [*Ca*], C2_08020C_A; *Neurospora crassa* [*Nc*], NCU02738; *Botrytis cinerea* [*Nc*], Bcin06g03400; *Aspergillus fumigatus* [*Af*], Afu3g08540). The position of the conserved five EF domains, each composed of two α-helices and one loop, was determined with the web server “PredictProtein” (https://predictprotein.org/). Disordered regions were predicted with the integrative protein classification web tool “InterPro” (https://www.ebi.ac.uk/interpro/). The protein schematics were illustrated with the “IBS 2.0” (https://ibs.renlab.org/#/home) online tool. **B:** Percentages of identity and similarity in between the aa sequences of the disordered regions (left) and the PEF-hand regions (right) from the human and fungal proteins presented in panel A. The values were calculated from aa alignments of these regions using the default setting of the online tool “Ident and Sim” at the Sequence Manipulation Suite (https://www.bioinformatics.org/sms2/index.html). All aa alignments were generated with the T-Coffee web server (https://tcoffee.crg.eu/apps/tcoffee/do:regular). n/a: not applicable.

**Fig S2:**
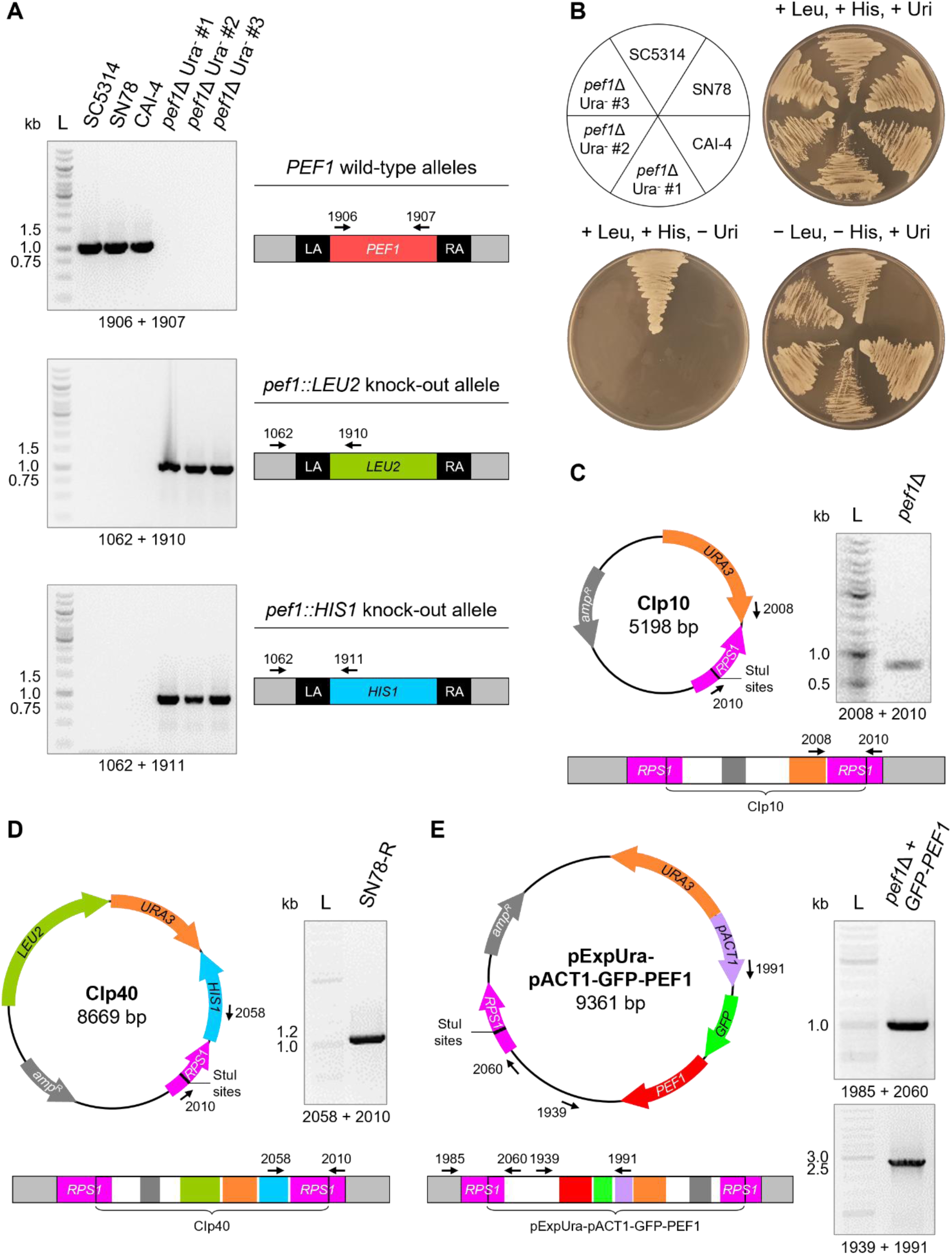
Construction and verification of the *pef1*Δ mutant and the GFP-Pef1 reporter strain. **A:** PCR analysis (left images) of three homozygous *pef1*Δ Ura^-^ mutants (MS-01 to -03), their parental strain SN78, and two control strains (SC5314, CAI-4). The mutants were generated through successive replacement of both *PEF1* alleles in SN78 by gene deletion cassettes containing the *LEU2* and *HIS1* selectable markers, respectively. The indicated primer pairs (numbered arrows) are specific for the wild-type or knock-out alleles (schematics to the right; not to scale). **B:** Growth of the strains from panel A on plates containing SD medium with or without the indicated supplements (Leu: leucine, His: histidine, Uri: uridine). The images of the plates were captured after 2 d of incubation at 30°C. **C:** PCR analysis (right image) of the prototrophic *pef1*Δ mutant (MW-Ca27) with the indicated primer pair to verify the integration of the *URA3* marker at the *RPS1* locus (linear schematic below) after transformation of the *pef1*Δ Ura^-^ mutant (MS-01) with the StuI-linearized *Candida* integrating plasmid, CIp10 (circular schematic at the left). **D:** PCR analysis of prototrophic parental strain, SN78-R (MW-Ca81), with the indicated primer pair. This control strain was generated through simultaneous re-integration of the *LEU2*, *URA3* and *HIS1* markers at the *RPS1* locus of SN78 via transformation with the StuI-linearized plasmid CIp40. **E:** PCR analysis of the *pef1*Δ strain complemented with the *GFP-PEF1* construct (MW-Ca58) using the indicated primer pairs. Complementation of the *pef1*Δ Ura^-^ mutant (MS-01) was achieved by transformation with the StuI-linearized vector pExpUra-pACT1-GFP-PEF1, which mediates constitutive expression of *GFP-PEF1* at the *RPS1* locus and also restores uracil prototrophy.

**Fig S3:**
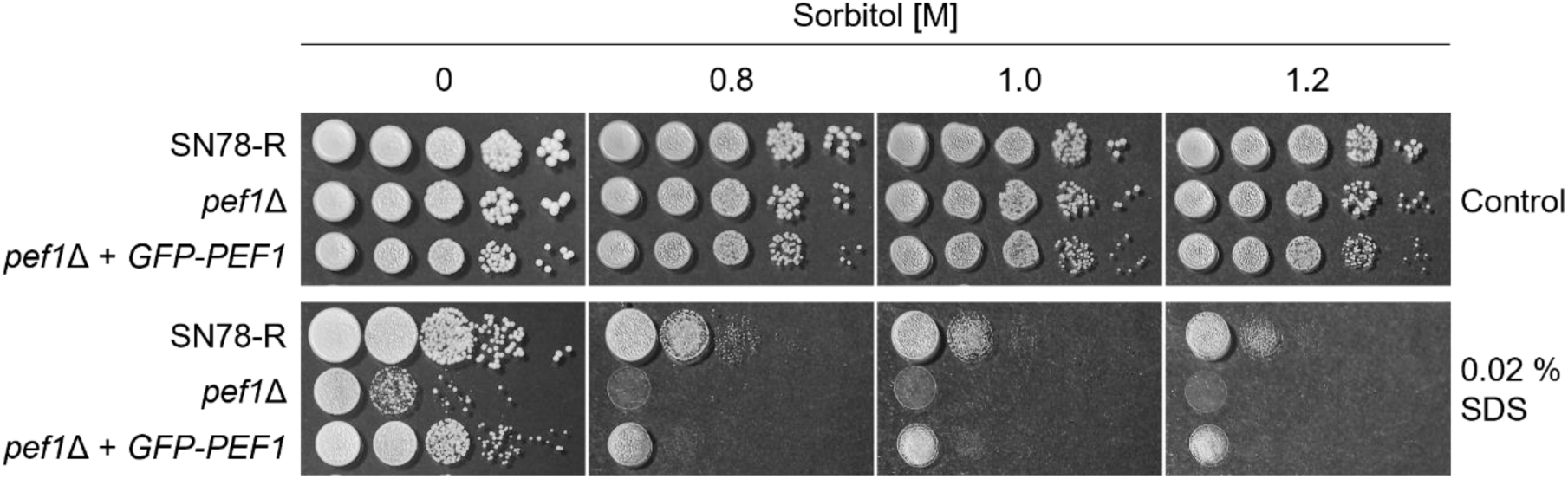
**Sorbitol does not protect the *pef1*Δ mutant from SDS.** Growth of colonies from 10-fold serial dilutions of yeast cells of SN78-R (MW-Ca81), the *pef1*Δ mutant (MW-Ca27) and the complemented mutant (MW-Ca58) on YPD agar plates supplemented with SDS and/or sorbitol. The images of the colonies were captured after incubation for 2 d at 37°C.

**Fig S4:**
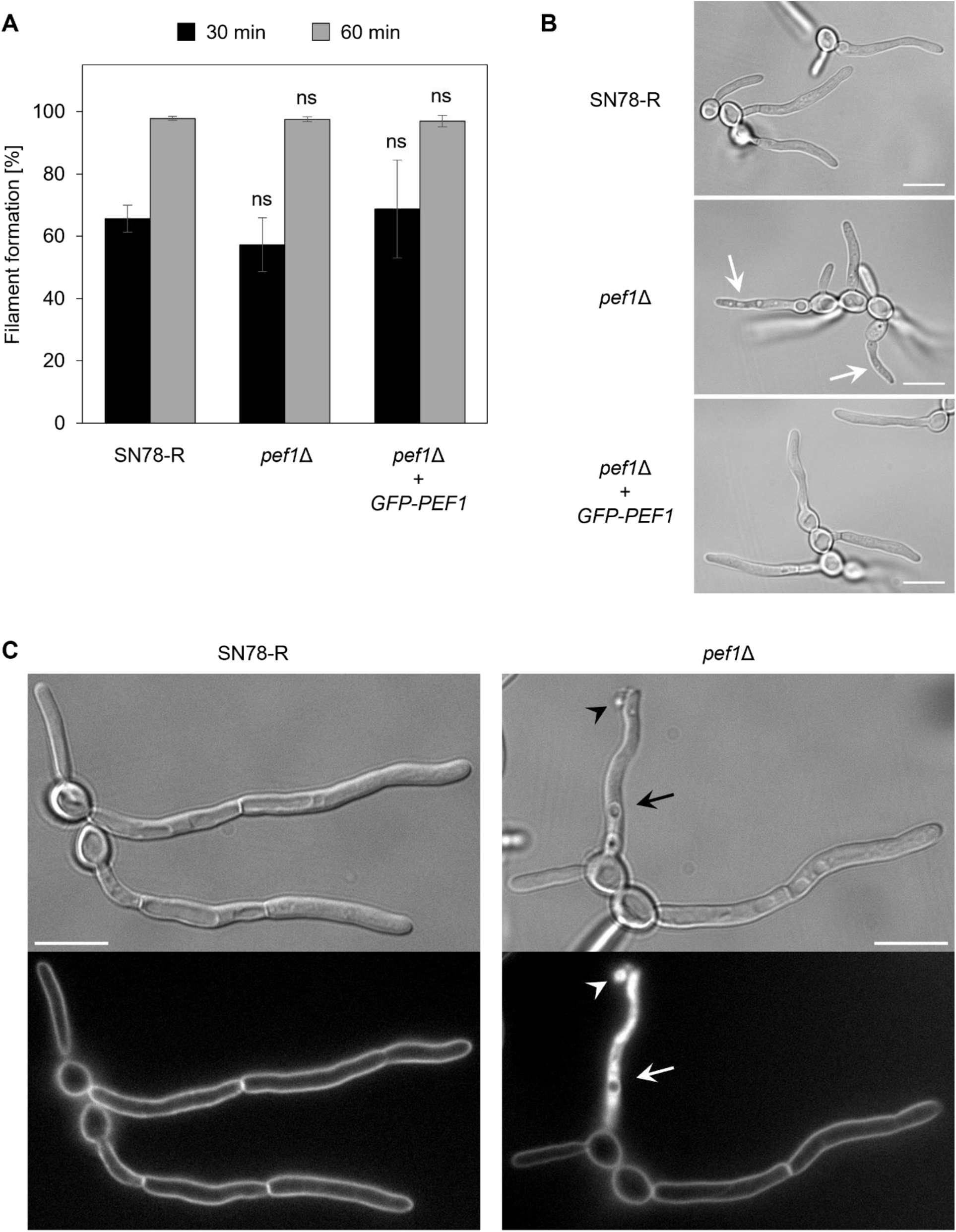
Pef1 is dispensable for the induction of filamentation but required for maintaining hyphal integrity. **A:** Quantification of the formation of filaments in the control strain SN78-R (MW-Ca81), the *pef1*Δ mutant (MW-Ca27) and the complemented mutant (MW-Ca58). Hyphae were induced by incubating yeast cells at 37°C in 20 % FBS. The percentage of filaments formed after 30 and 60 min was determined from three technical replicates per strain and time point. Mean values (black and gray bars) and errors bars (Std Dev) were statistically compared with each other by one-way ANOVA analysis with Tukey’s correction for multiple comparisons (ns, not significant). **B:** Bright-field images of filaments of the strains from panel A after incubation for 2 h at 37°C in 20 % FBS. The hyphae of the *pef1*Δ mutant show signs of stress in the form of vacuolization (arrows). Scale bars: 10 μm. **C:** FM4-64 staining of hyphae from the indicated strains grown at 37°C in 20 % FBS. Fluorescence images (bottom) show plasma membrane staining in all filaments, whereas accumulation of the dye is only visible inside one of the mutant hyphae (white arrow). That filament appears vacuolized in the DIC image above (black arrow) and shows leakage of cellular content (back arrowhead) that is stained with FM4-64 (white arrowhead) is visible. Scale bars: 10 μm.

**Fig S5:**
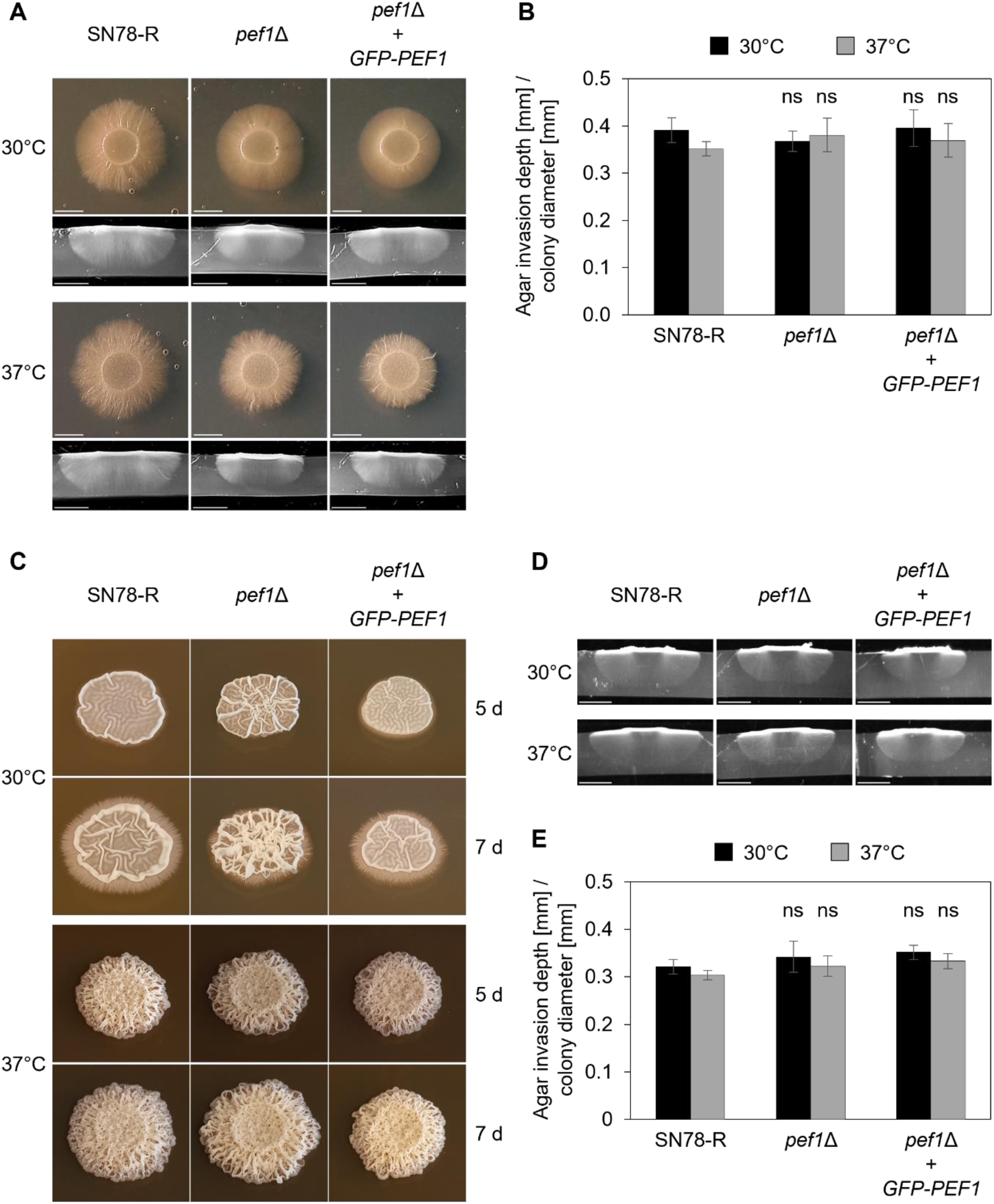
Loss of Pef1 does not impair invasive filamentous growth of *C. albicans* on solid media. **A:** Growth of SN78-R (MW-Ca81), the *pef1*Δ mutant (MW-Ca27) and the complemented strain (MW-Ca58) on solid serum medium (10 % FBS, 2 % agar) at 30°C and 37°C. The top images show the filamentation of the colonies after 7 d of incubation. Representative cross-sections prepared from the colonies are shown in the images below. Scale bars: 5 mm. **B:** Quantification of the ratio between the average depth of agar invasion and the average diameter of colonies grown on serum agar as shown in panel A. Mean values and error bars (Std Dev) derived from two technical replicates were statistically analyzed by one-way ANOVA analysis with Tukey’s correction for multiple comparisons (ns, not significant). **C:** Representative colonies of the same strains as presented in panel A grown on solid Spider medium after 5 d and 7 d of incubation at 30°C and 37°C. **D:** Images of representative cross-sections prepared from colonies grown on Spider agar as shown in panel C after 7 d of incubation at 30°C and 37°C. Scale bars: 5 mm. **E:** Quantification of the ratio between the average depth of agar invasion and the average diameter of colonies grown on Spider agar as shown in C and D. Mean values and error bars (Std Dev) derived from four technical replicates were statistically analyzed by one-way ANOVA analysis with Tukey’s correction for multiple comparisons (ns, not significant).

**Fig S6:**
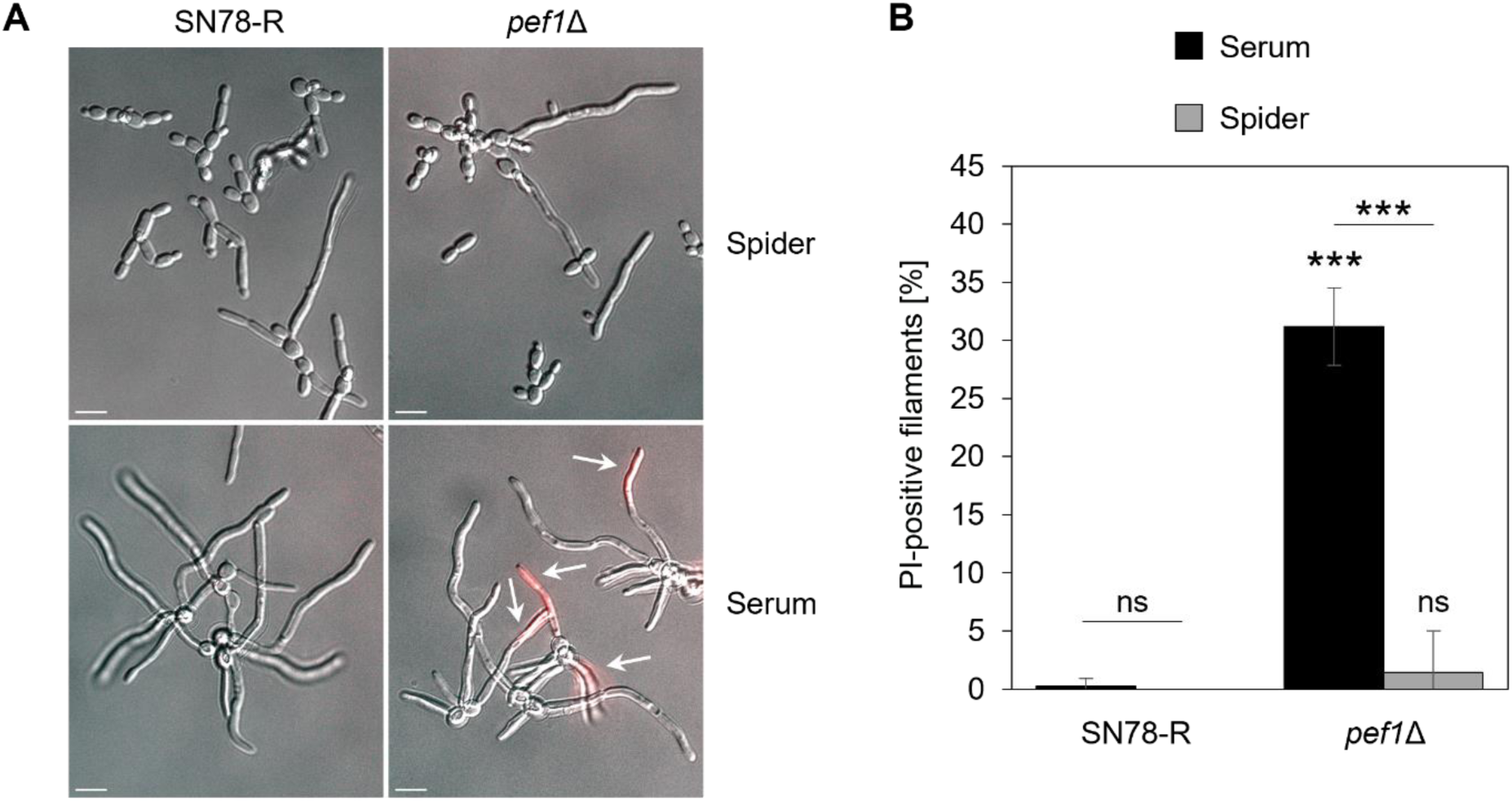
Loss of Pef1 does not impair hyphal integrity in liquid Spider medium. **A:** Filamentation of the *pef1*Δ mutant (MW-Ca27) and its control strain (MW-Ca81) after incubation at 37°C for 3-4 h in liquid Spider medium or 20 % FBS (serum). Images show overlays from DIC and fluorescence microscopy of cells stained with red-fluorescent propidium iodide (PI). **B:** Quantification of the percentage of PI staining in filaments from the culture conditions shown in panel A. The black and gray bars represent mean values with errors (Std Dev) from 5-6 technical replicates obtained from two independent experiments per strain and medium. Statistically significant differences (***, p < 0.001; ns, not significant) were assessed by one-way ANOVA analysis with Tukey’s correction for multiple comparisons.

**Fig S7:**
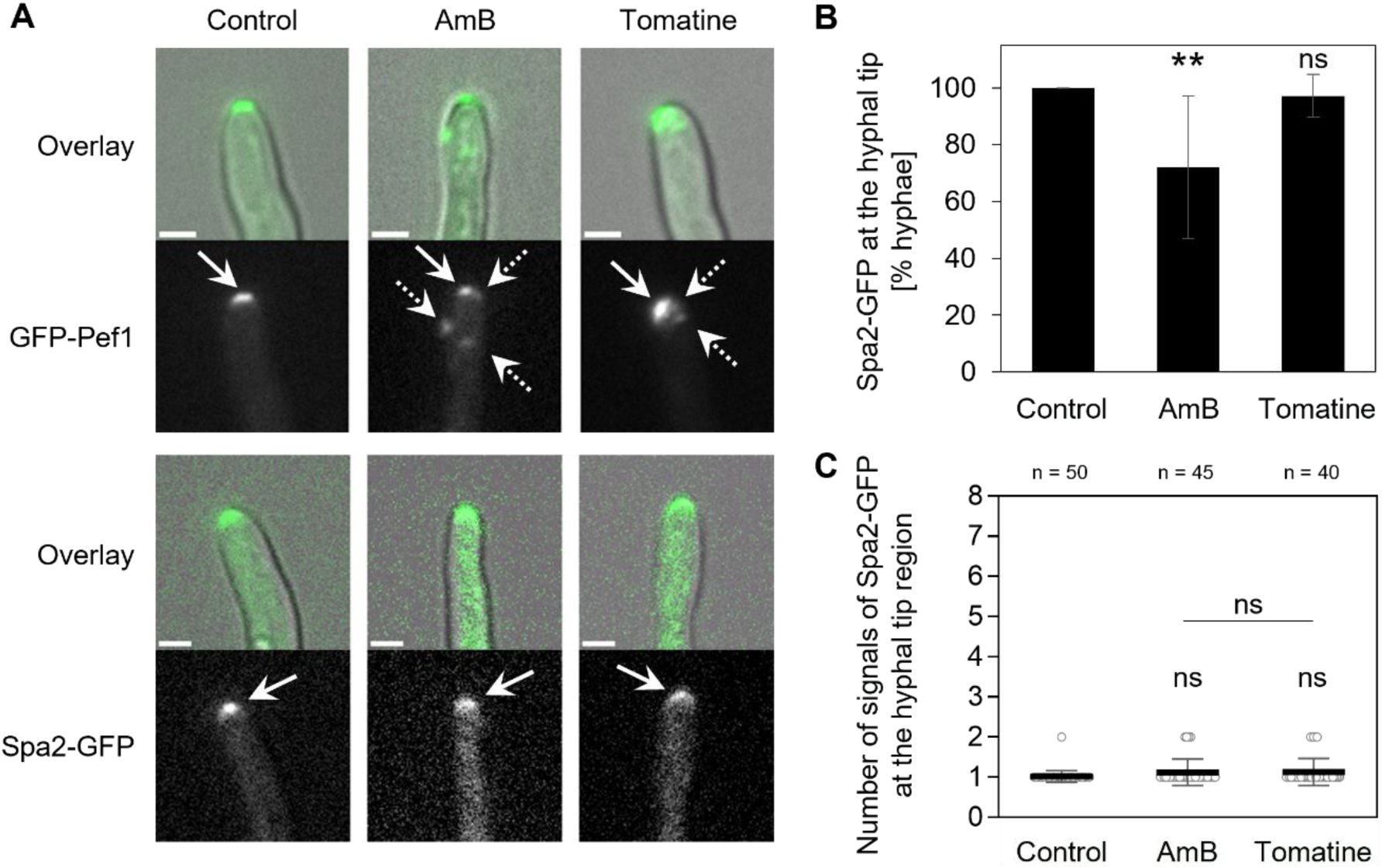
Membrane stress does not alter the localization pattern of the polarisome marker protein Spa2. **A:** Fluorescence microscopy comparing the localization patterns of GFP-Pef1 (in the reporter strain MW-Ca58) and Spa2-GFP (strain WYZ6) in hyphae grown for 3 h at 37°C in 20 % FBS prior to treatment with 2 µg/mL AmB or 50 µg/mL tomatine for up to 15 min. In addition to the localization of GFP-Pef1 at the hyphal tips (solid arrows), other punctate signals of the fluorescent protein accumulate at the hyphal tip region (dotted arrows) in response to the membrane-disrupting antifungal compounds (see also Fig. 4). In contrast, antifungal treatment does not alter the localization pattern of Spa2-GFP at the hyphal tip region, which is comparable to the one observed for GFP-Pef1 in the absence of membrane stress (solid arrows). Overlay: merged DIC and fluorescence images. AmB: Amphotericin B. Scale bars: 2 μm. **B:** Quantification of the localization of Spa2-GFP at the hyphal tip of filaments as shown in A. The black bars show the mean values with errors (Std Dev) from 8-9 technical replicates per condition with 3-12 filaments per replicate. **C:** Quantitative analysis of the signal distribution of Spa2-GFP at the hyphal tip of the filaments from panel B in the absence and presence of AmB or tomatine. The dot plot shows the number of distinct fluorescent signals observed at the hyphal tip region of each filament (open circles) as well as the mean value and error bars (Std Dev) from the total number (n) of filaments analyzed per condition. Statistically significant differences in B and C were assessed by one-way ANOVA analysis with Tukey’s correction for multiple comparisons (**, p < 0.01; ns, not significant).

**Fig S8:**
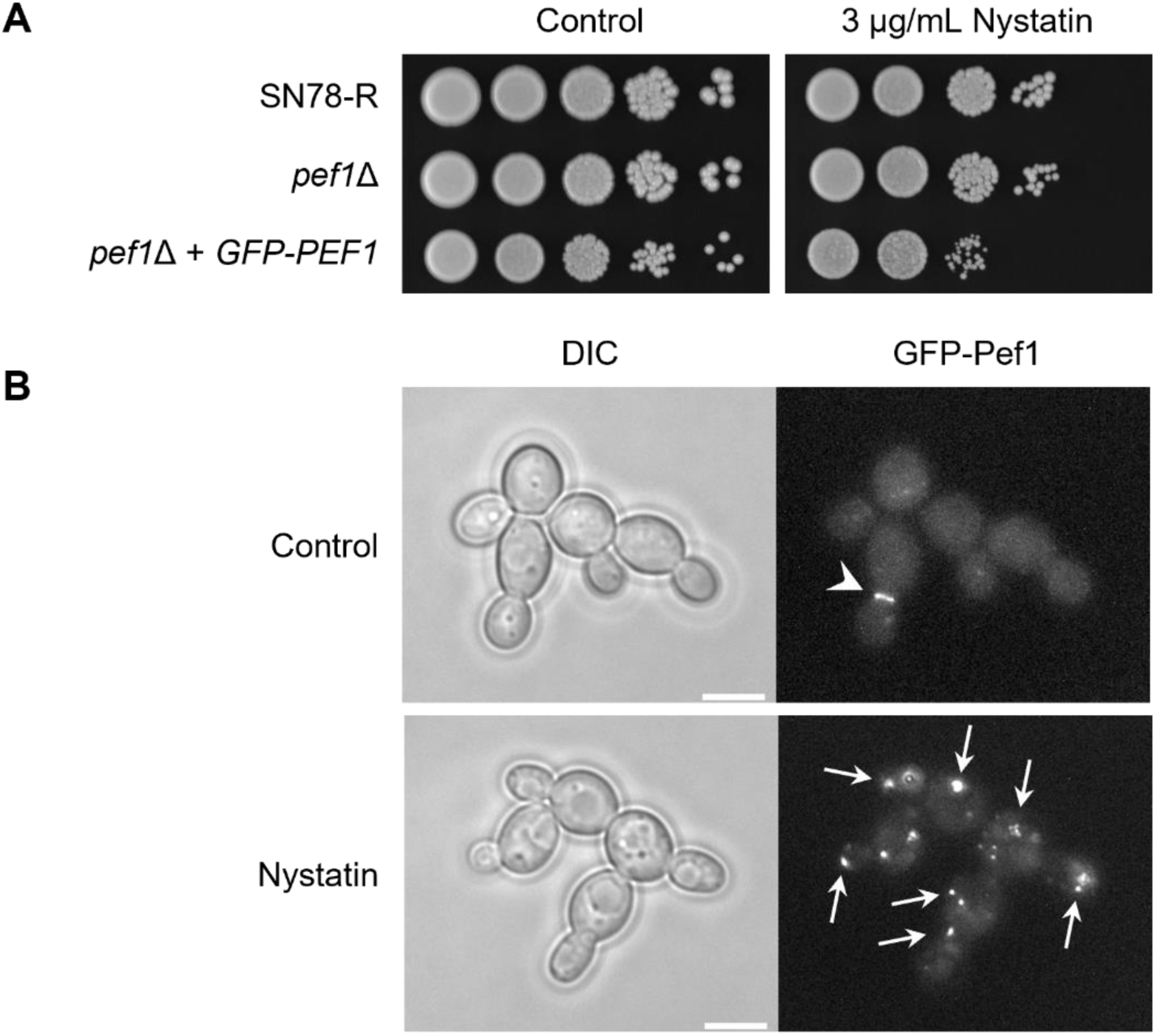
Pef1 alters its localization pattern during the exposure of yeast cells to nystatin but is dispensable for the adaptation to this polyene. **A:** Colonies grown from 10-fold serial dilutions of yeast cells of SN78-R (MW-Ca81), the *pef1*Δ mutant (MW-Ca27) and the complemented strain (MW-Ca58) were spotted on solid YPD medium with and without nystatin. The images of the plates were captured after incubation for 2 d at 37°C. **B:** Localization of GFP-Pef1 in yeast cells of the reporter strain (MW-Ca58) exposed to nystatin. Cells from overnight cultures were grown for 6 h at 30°C in fresh YPD broth and washed in 0.9% saline prior to treatment with this polyene. Images were captured after about 10 min of incubation in the presence of 20 μg/mL nystatin or 0.2 % DMSO (control). Arrowhead: normal localization of GFP-Pef1 at the bud neck of dividing control cells; arrows: punctate accumulation of GFP-Pef1 at the cell periphery and at intracellular sites after antifungal treatment. Scale bars: 5 μm.

**Fig S9:**
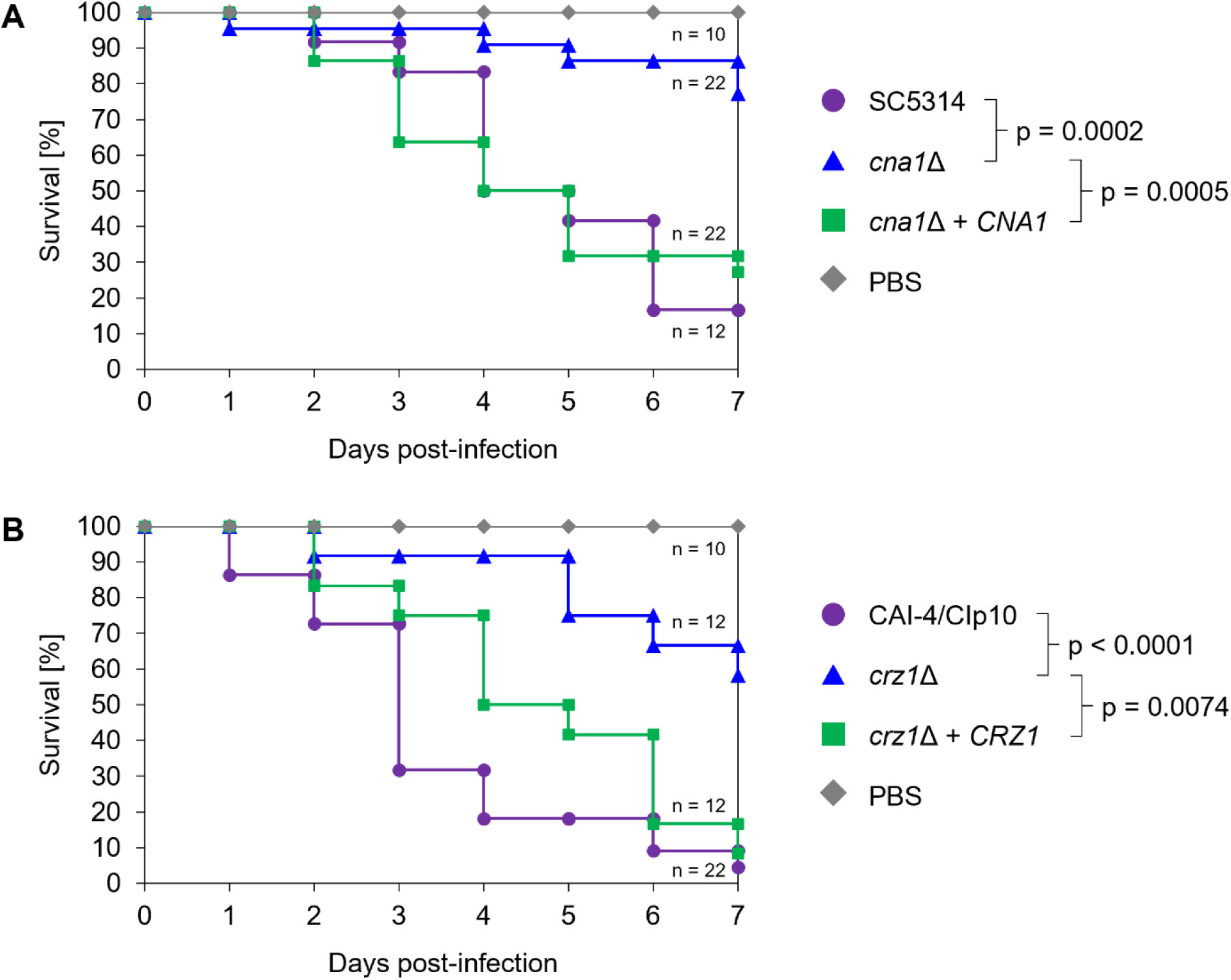
Loss of the Ca^2+^/calcineurin signaling proteins Cna1 and Crz1 attenuates virulence of *C. albicans* in an insect larvae infection model. Survival plots of *G. mellonella* larvae infected with yeast cells of two different sets of strains of *C. albicans*: **A**, the wild-type strain SC5314, the *cna1*Δ mutant (SCCMP1M4) and the complemented *cna1*Δ mutant (SCCMP1MK2); **B**, the control strain CAI-4/CIp10, the *crz1*Δ mutant (MKY380) and the complemented *crz1*Δ mutant (MKY381). After injection on day 0 with 1x10^6^ cells per larva or with sterile PBS as a control, all larvae were incubated for one week at 37°C and checked daily for survival. The plots represent pooled data from two independent experiments with the indicated total number (n) of larvae used per strain. Statistically significant differences were assessed by Log-rank analysis with the indicated p values.

